# Heterogeneity in the gene regulatory landscape of leiomyosarcoma

**DOI:** 10.1101/2022.04.13.488196

**Authors:** Tatiana Belova, Nicola Biondi, Ping-Han Hsieh, Pavlo Lutsik, Priya Chudasama, Marieke L. Kuijjer

## Abstract

Soft-tissue sarcomas are group of rare, tremendously heterogeneous, and highly aggressive malignancies. Characterizing inter-tumor heterogeneity is crucial for selecting suitable sarcoma therapy, as the presence of diverse molecular subgroups of patients can be associated with disease outcome or response to treatment. While cancer subtypes are often characterized by differences in gene expression, the mechanisms that drive these differences are generally unknown. We therefore set out to model the regulatory mechanisms driving sarcoma heterogeneity. We subtyped soft-tissue sarcomas based on patient-specific, genome-wide gene regulatory networks and found pronounced regulatory heterogeneity in leiomyosarcoma—one of the most common soft-tissue sarcomas subtypes that arises in smooth muscle tissue. To characterize this regulatory heterogeneity, we developed a new computational framework. This method, PORCUPINE, combines knowledge on biological pathways with permutation-based network analysis to identify pathways that exhibit significant regulatory heterogeneity across a patient population. We applied PORCUPINE to patient-specific leiomyosarcoma networks modeled on data from The Cancer Genome Atlas and validated our results in an independent dataset from the German Cancer Research Center. PORCUPINE identified 37 heterogeneously regulated pathways, including pathways that represent potential targets for treatment of subgroups of leiomyosarcoma patients, such as FGFR and CTLA4 inhibitory signaling. We validated the detected regulatory heterogeneity through analysis of networks and chromatin states in leiomyosarcoma cell lines. In addition, we showed that the heterogeneity identified with PORCUPINE is not associated with methylation profiles or clinical features, thereby suggesting an independent mechanism of patient heterogeneity driven by the complex landscape of gene regulatory interactions.

## I. INTRODUCTION

Soft-tissue sarcomas are a group of rare and highly aggressive malignancies. While they account for less than 1% of all malignant tumors, soft-tissue sarcomas are a tremendously heterogeneous group of tumors and include more than 150 different histological subtypes [1]. Partly because of this heterogeneity, significant challenges exist in the management of soft-tissue sarcomas. Most soft-tissue sarcomas are treated similarly in the clinic, regardless of their site of origin, with surgery with or without radiotherapy as the main treatment for localized disease [2]. Several clinical trials have been conducted in soft-tissue sarcomas. However, until recently such trials included patients with many different histological subtypes in the same cohort, causing difficulties to conclude on the efficacy of these therapies in the individual subtypes. Differences in clinical response among soft-tissue sarcoma subtypes led to newer studies that only enrolled patients of certain histological subtypes, which have shown to result in better response and disease control [3].

Over the past years it has become evident that treatments tailored to a single patient, or group of patients belonging to a specific molecular subtype of cancer, can result in major improvements in cancer outcomes [4]. For example, characterizing inter-patient molecular tumor heterogeneity was shown to be crucial for selecting the most efficient cancer therapy, and the presence of diverse molecular subtypes can predict patient survival in breast cancer [5] and relapse or resistance to treatment in melanoma [6]. Therefore, it is clear that the integration of personalized medicine into cancer treatment strategies requires extensive knowledge of inter-patient variability. Patients can, for example, be grouped into molecular subtypes based on “omics” data, such as gene expression, microRNA, DNA methylation, somatic mutations, or proteomic profiles.

The molecular landscape of soft-tissue sarcomas has been characterized in several studies [7–10]. The Cancer Genome Atlas (TCGA) sarcoma project, one of the largest sarcoma sequencing projects to-date, performed a comprehensive and integrated analysis of 206 adult soft-tissue sarcomas, represented by six major subtypes, and showed that sarcomas vary greatly at the genetic, epigenetic, and transcriptomic levels [7]. More recently, some histological subtypes of soft-tissue sarcomas were further delineated into molecular subgroups according to their genomic and transcriptomic profiles. For example, Guo *et al.*, characterized three molecular subtypes of leiomyosarcoma (LMS)—one of the most common subtypes of soft-tissue sarcomas—based on transcriptomic data. One of these subtypes was over-represented by uterine leiomyosarcoma, while the other two were over-represented by extra-uterine sites. While these subtypes were not associated with tumor grade, they were somewhat related to patient survival [11]. However, the causative regulatory mechanisms that distinguish these subtypes are not fully understood and the impact of molecular profiling of soft-tissue sarcomas on patient outcomes has been limited.

Through the modeling of interactions between transcription factors (TFs) or other regulators and their potential target genes, gene regulatory networks offer an in-depth view on the mechanisms that drive gene expression [12], and thus could help gain greater insight into disease mechanisms. Various integrative methods have been developed to model such networks genome-wide. One such method is PANDA, which integrates putative TF-DNA binding with protein-protein interactions and target gene co-expression to infer a regulatory network for a specific condition [13]. Recently, we developed an algorithm that can be combined with condition-specific network models estimated with e.g. PANDA to infer patient-specific regulatory networks (LIONESS [14]). These patient-specific network models have been instrumental in capturing sex differences in gene regulation in healthy tissues [15] and colon cancer [16], as well as in identifying regulatory interactions associated with glioblastoma survival [17].

In this work, we demonstrate that analysis of heterogeneity among patient-specific gene regulatory networks can facilitate stratification of soft-tissue sarcoma patients into novel regulatory subtypes and identification of the regulatory programs that drive such heterogeneity. We identified a high level of regulatory heterogeneity in leiomyosarcoma. To characterize this heterogeneity, we present a new computational approach, PORCUPINE (Principal Components Analysis to <u>O</u>btain <u>R</u>egulatory <u>C</u>ontributions <u>U</u>sing <u>P</u>athway-based Interpretation of <u>N</u>etwork <u>E</u>stimates), to detect statistically significant, key regulatory pathways that drive regulatory heterogeneity among patients.

We applied PORCUPINE to 80 genome-wide leiomyosarcoma regulatory networks, which we modeled on data from TCGA (referred to below as TCGA-LMS). We validated the pathways detected by PORCUPINE in an independent dataset consisting of 37 leiomyosarcoma available from the study by Chudasama *et al.* (referred to below as DKFZ-LMS) [18]. We found high concordance in regulatory heterogeneity in both cohorts, identifying 37 shared heterogeneously regulated pathways. These included pathways that play a known role in leiomyosarcoma biology and pathways that have not been described before in the disease. Newly identified pathways include FGFR signaling and CTLA4 inhibitory signaling and represent potential targets for treatment of subgroups of leiomyosarcoma patients. We validated the detected regulatory heterogeneity through analysis of networks and chromatin states in leiomyosarcoma cell lines. Moreover, we show that the heterogeneity identified with PORCUPINE is not associated with methylation profiles or clinical features, thereby suggesting an independent mechanism of patient heterogeneity driven by the complex landscape of gene regulatory interactions.

## MATERIALS AND METHODS

### Gene expression data preprocessing

We downloaded expression data for all TCGA cases using the “recount” package in R [19]. The transcriptome data for 37 leiomyosarcoma cases obtained from the German Cancer Research Center (DKFZ) was preprocessed by the Omics IT and Data Management Core Facility (DKFZ ODCF) using the One Touch Pipeline [20]. We performed batch correction on the raw expression counts of the set of 206 TCGA soft-tissue sarcomas and the 37 DKFZ-LMS samples together, using the “Combat-seq” package in Bioconductor [21]. We then combined Combat-seq-adjusted counts with the raw expression counts of the remaining TCGA samples and performed smooth quantile normalization using “qsmooth” package in Bioconductor to preserve global differences in gene expression between the different cancer types [22], specifying each cancer type as a separate group level. Samples of 206 TCGA soft-tissue sarcomas and 37 DKFZ-LMS samples were specified as the same “soft-tissue sarcoma” group level.

### Construction of individual patient gene regulatory networks

We used the MATLAB version of the PANDA network reconstruction algorithm (available in the net-Zoo repository https://github.com/netZoo/netZooM) to estimate an “aggregate” gene regulatory network, based on a total of 11,321 samples, 17,899 genes, and 623 TFs. These samples included 206 TCGA and 37 DKFZ soft-tissue sarcomas—the remaining samples represented other cancer types available in TCGA. We used the entire TCGA dataset to build the aggregate network, as we previously found that LIONESS’ estimates of single-sample edges are more robust when including a large, heterogeneous background of samples [14].

PANDA builds an aggregate network by incorporating three types of data—a “prior” regulatory network, which is based on a TF motif scan to identify putative regulatory interactions between TFs and their target genes, protein-protein (PPI) interactions between TFs, and target gene expression data. The aggregate network modeled by PANDA consists of weighted edges between each TF-target gene pair. These edge weights reflect the strength of the inferred regulatory relationship.

The prior gene regulatory network was generated using a set of TF motifs obtained from the Catalogue of Inferred Sequence Binding Preferences (CIS-BP) [23], as described by Sonawane *et al.*, 2017 [24]. These motifs were scanned to promoters as described previously [25]. The prior network was intersected with the expression data to include genes and TFs with available expression data and at least one significant promoter hit. This resulted in initial map representing potential regulatory interactions between 623 TFs and 17,899 target genes. An initial protein-protein network was estimated between all TFs from motif prior map using interaction scores from StringDb v10 [26], which were scaled to be within a range of [0,1], where self-interactions were set equal to one, as described previously [24]. To reconstruct patient-specific gene regulatory networks, we applied the LIONESS equation in MATLAB (available in the netZoo repository https://github.com/netZoo/netZooM).

### UMAP visualization

To visualize the clustering distribution of the 206 TCGA soft-tissue sarcoma patient-specific gene regulatory networks, we applied dimensionality reduction with Uniform Manifold Approximation and Projection (UMAP), using the “uwot” package in R 3.6.1, setting the number of nearest neighbours to 20. We performed UMAP on the matrix of gene targeting scores obtained from the 206 individual sarcoma networks. Gene targeting scores are defined as the sum of all edge weights pointing to a gene and represent the amount of regulation a gene receives from the entire set of TFs available in a network [27]. These scores have previously been used to identify gene regulatory differences in various studies [16, 17, 27]. We visualized the results in two-dimensional UMAP space. To identify clusters in the data, we used the HDBSCAN clustering algorithm on the UMAP coordinates from the first two embeddings [28], with the parameter “minPts” set to five.

### Identifying regulatory heterogeneity using PORCUPINE

To capture inter-patient heterogeneity (referred to below as “heterogeneity”) at the gene regulatory level, we developed a computational framework, which we call PORCUPINE. PORCUPINE is a Principal Components Analysis (PCA)-based approach that can be used to identify key pathways that drive heterogeneity among individuals in a dataset. It determines whether a specific set of variables—for example a set of genes in a specific pathway—have coordinated variability in their regulation.

PORCUPINE uses as input individual patient networks, for example networks modeled using PANDA and LIONESS, as well as a.gmt file (in MSigDb file format [29]) that includes biological pathways and the genes belonging to them. For each pathway, it extracts all edges connected to the genes belonging to that pathway and scales each edge across individuals. It then performs a PCA analysis on these edge weights, as well as on a null background that is based on random pathways. For the randomization (permutation), PORCUPINE creates a set of 1000 gene sets equal in size to the pathway of interest, where genes are randomly selected from all genes present in the.gmt file. The edges connected to these genes are then extracted. The amount of variance explained by the first principal component (PC1) in the pathway of interest is then compared to the amount of variance explained by PC1 in the random (permuted) data.

To identify significant pathways, PORCUPINE applies a one-tailed t-test and calculates the effect size (ES). The latter is calculated as the difference between the variance explained by PC1 of the pathway of interest and the mean of the variance explained by PC1 corresponding to the random sets of pathways, divided by standard deviation of the variance explained by PC1 in the random sets using the cohensD function in the “lsr” package in R. P-values are adjusted for multiple testing with the Benjamini-Hochberg method [30] and significant pathways are returned based on user-defined thresholds of adjusted p-value and effect size. We developed PORCUPINE as R package and it is available as open-source code on GitHub (https://github.com/kuijjerlab/PORCUPINE).

We applied PORCUPINE to TCGA and DKFZ leiomyosarcoma data using Reactome pathways v7.1 from MSigDb, excluding pathways that consisted of more than 200 genes. Pathways with adjusted p-value less than 0.01 and effect size >=2 were reported as significant. As the number of genes in each pathway is different, we investigated whether the obtained results were biased towards pathways of smaller size. To test this, we split pathways in four groups based on their size, namely pathways containing less than 50, 50-100, 100-150, 150-200 genes. We then calculated the proportions of these groups among Reactome pathways and among the set of deregulated pathways identified in the TCGA-LMS and DKFZ-LMS datasets.

### Clustering of pathways and identification of redundant aspects of gene regulatory heterogeneity

To investigate potential redundant patterns of heterogeneity captured by pathways identified with PORCUPINE, we computed the Pearson correlation coefficient for every pair of individuals for each pathway, based on the individual’s TF-target edge weights in that pathway. We then combined pathway-level inter-individual correlations into a matrix for all pathways and performed clustering, visualizing the results using the “ComplexHeatmap” package in R. Additionally, to identify pathways with overlapping genes, we computed the Jaccard similarity between pairs of pathways.

### Identification of top ranked target genes and transcription factors

To identify those genes and TFs that contribute most to the pathway’s significance, we extracted the edge loadings of the first principal component (referred to below as the “edge contribution score”). Because the sum of the squares of all edge contribution scores for an individual principal component must be one, we calculated the expected edge contribution score, assuming that all edges contributed equally to that principal component. Edges with a contribution score > 1.5× the expected score were regarded as important contributors to that principal component. To identify TFs with many co-regulated genes, we then grouped TFs corresponding to these top edges according to the number of their targets.

### Association of the significant pathways with clinical phenotypes

To investigate whether the heterogeneity captured by each pathway was associated with clinical features, we performed an association analysis of the coordinates of patients on the first principal component in each pathway (referred to below as the “pathway-based patient heterogeneity score”) with the clinical data available for these patients. Clinical features for TCGA leiomyosarcoma patients were obtained using the “TCGAbiolinks” package from Bioconductor [31]. Clinical information for 37 DKFZ patients was obtained from the study by Chudasama *et al.* [18]. Since the clinical attributes represent a mix of categorical and numerical features, we applied Kruskal-Wallis and Pearson correlation tests for categorical and numerical features, respectively. We corrected p-values for multiple testing using the Benjamini-Hochberg approach and applied a threshold of 0.05 to identify significant associations.

In order to determine whether any of the identified pathways were associated with patient survival, we used the first principal component from each pathway in a Cox regression model to predict patient survival.

### Association of the significant pathways with pathway-based mutation profiles

We downloaded and preprocessed leiomyosarcoma mutation data as previously described in Kuijjer *et al.* [32]. We used the SAMBAR algorithm [32] to obtain patient-specific pathway mutation scores for TCGA-LMS patients. Among 1,455 pathways, 954 pathways had mutation scores larger than zero in the TCGA-LMS dataset. To assess the association between pathways identified with PORCUPINE and these pathways mutation scores, we used a Kruskal Wallis test, comparing the pathway-based patient heterogeneity scores on the first principal component between two groups, i.e. mutated vs not mutated, for each mutated pathway. We used FDR <0.05 as threshold for reporting significant differences between the groups.

### Association of the identified pathways with overall methylation profiles

DNA methylation data measured on the Illumina Infinium Human Methylation 450 BeadChip platform were downloaded for all sarcoma patients available in TCGA using the Bioconductor “TCGA biolinks” package in R. We downloaded raw methylation IDAT files and performed preprocessing and normalization with subset-quantile within array normalization (SWAN) using Bioconductor package “minfi.” [33]. We calculated overall methylation profiles for each individual by using the mean value across all probes. We then correlated these values to the pathway-based patient heterogeneity scores in each pathway. Associations with FDR <0.05 were considered significant.

### Validation of the pathways in healthy tissues

We obtained patient-specific regulatory networks for healthy smooth-muscle–derived tissues, represented by esophageal muscularis and uterus from the Genotype-Tissue Expression (GTEx) project, through the GRAND database of gene regulatory network models [34]. In total, 283 and 90 patient-specific networks were available for esophageal muscularis and uterus, respectively. We applied PORCUPINE to evaluate gene regulatory heterogeneity among the individuals in the merged set of 373 networks.

### Construction of gene regulatory networks for leiomyosarcoma cell lines

RNA-seq counts were obtained for four leiomyosarcoma cell lines, including SK-UT-1, SK-UT1-B, MES-SA and SK-LMS-1 from the study by Chudasama *et al.* [18]. To integrate these data with the patient samples, we performed batch correction on the raw expression counts of the set of 206 TCGA soft-tissue sarcomas and the 37 DKFZ-LMS samples and the 4 cell lines together, using the “Combatseq” package in Bioconductor. Following that, we combined the Combatseq-adjusted counts with the raw expression counts of the remaining TCGA samples and used “qsmooth” normalization to obtain normalized counts. Individual networks for cell lines were then modeled using PANDA and LIONESS as described above.

### Generation and processing of ATAC-seq data

A complete list of all reagents, buffer solutions, and DNA barcode primer sequences is described in Supplementary File 1. ATAC-Seq libraries for SK-UT-1, SK-UT-1B, MES-SA, and SK-LMS-1 were prepared in triplicate according to the Omni-ATAC protocol [35] with minor modifications. Briefly, 50,000 cells per replicate were collected by centrifugation at 500 x g for 5 min at 4°C. Cell pellets were resuspended in 50 *μ*L ice-cold lysis buffer A and incubated on ice for 3 min, after which 500 *μ*L ice-cold lysis buffer B were added. Cell nuclei were pelleted by centrifugation at 500 x g for 10 min at 4°C. The supernatant was removed carefully and nuclei pellets were resuspended in 47.5 *μ*L ice-cold transposition buffer and 2.5 *μ*L Tagment DNA TDE1 enzyme (Illumina). The transposition mix was incubated at 37°C for 30 mins at 1000 rpm. After adding 20 *μ*L 5 M guanidinium thiocyanate, tagmented DNA was purified using Agencourt AMPure XP magnetic beads (Beckman Coulter).

Sequencing libraries were generated via qPCR by mixing purified tagmented DNA with 25 *μ*L 2X NEB-Next High-Fidelity PCR Master Mix (NEB), 2.5 *μ*L Tn5mCP1n forward primer, 2.5 *μ*L Tn5mCBar reverse primer, and 0.5 *μ*L 100X SYBER Green I (Invitrogen). The following PCR program was implemented: 1 cycle of 72°C for 5 min, 1 cycle of 98°C 30 sec, and 10 cycles of 98°C for 10 sec, 63°C for 30 sec, 72°C for 30 sec. Following two-sided size selection with 0.5× and 1.4× of Agencourt AMPure XP magnetic beads, library concentration and fragment distribution were checked via the 2200 TapeStation System with the High Sensitivity D1000 ScreenTape/Reagents (Agilent Technologies).

Libraries were sequenced at the DKFZ Genomics and Proteomics Core Facility using the Illumina NextSeq 550 Paired-End 75 bp. Sequencing reads were processed using the CWL-based ATAC-Seq workflow available at https://github.com/CompEpigen/ATACseq_workflows [36, 37]. Peak calling on individual samples was performed with MACS2 with parameters – nomodel – keep dup all –broad –gsize 2736124973 –qvalue 0.05. We followed the DiffBind protocol to obtain a consensus read count matrix from MACS2 peak sets [38]. The ATAC-seq peaks were filtered using the ENCODE blacklist [39] and only the peaks present at least in any two samples were included in the analysis. Peaks were annotated to nearest gene using the “annotatePeak” function in “ChIPseeker” package in R. To identify differentially accessible regions between different cell lines we used the raw read count matrix in DESeq2 [40]. For this, only genomic regions that were annotated as promoter regions based on the annotatePeak calls were considered. If several promoters were mapped to the same gene, a mean of raw reads over those promoter regions was calculated. To obtain normalized ATAC counts for the comparison of peak accessibility at the promoters of the genes in heterogeneous pathways with random regions, we used library size normalization in DiffBind. If several promoters were mapped to the same gene, a mean of normalized reads over those promoter regions was calculated.

## RESULTS

### Pan-sarcoma clustering of patient-specific regulatory networks

We set out to investigate the regulatory processes that drive heterogeneity in soft-tissue sarcomas. We started by modeling genome-wide, patient-specific gene regulatory networks for 206 TCGA soft-tissue sarcoma patients using two computational algorithms, PANDA and LIONESS (Figure 1). These patient-specific networks include information on likelihoods of regulatory interactions (represented as edge weights) between 623 TFs and 17,899 target genes. To explore and visualize patient heterogeneity based on their regulatory landscapes, we first calculated gene targeting scores in these networks (see Methods), and then used Uniform Manifold Approximation and Projection (UMAP) for visualization. To determine whether regulatory profiles cluster differently than expression data, we also performed UMAP on the expression data (Figure 2).

**Figure 1.**
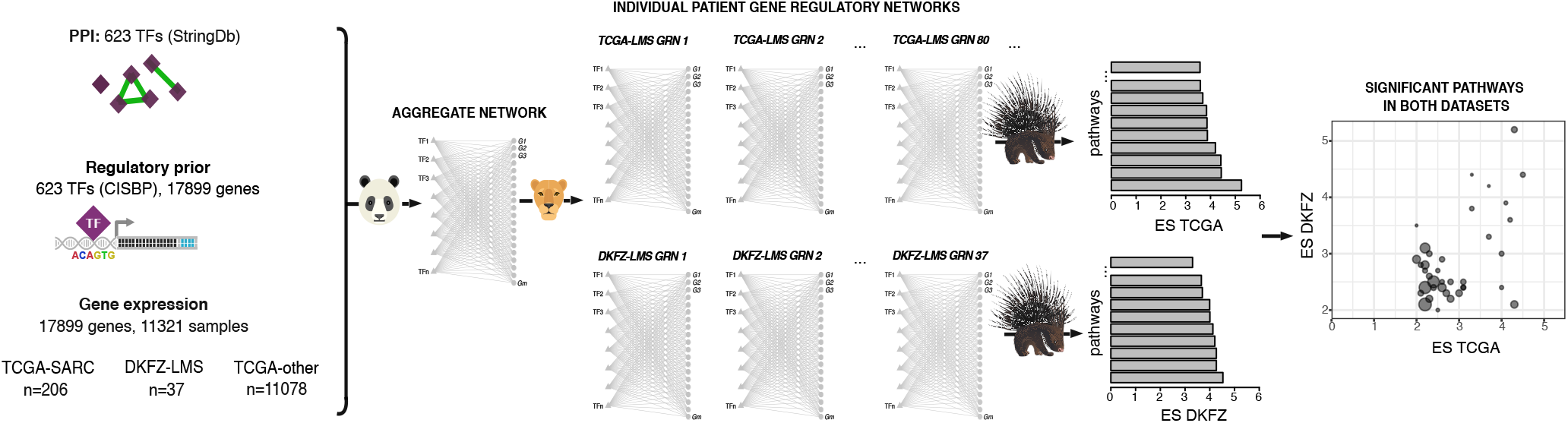
Schematic overview of the study. We modeled individual patient gene regulatory networks for leiomyosarcoma patients from two datasets (TCGA and DKFZ) with PANDA and LIONESS, integrating information on protein-protein interactions (PPI) between transcription factors (TF), prior information on TF-DNA motif binding, and gene expression data. We then developed and applied a new computational comparative network analysis tool (PORCUPINE) to identify significant pathways that capture heterogeneity in gene regulation across these datasets.

**Figure 2.**
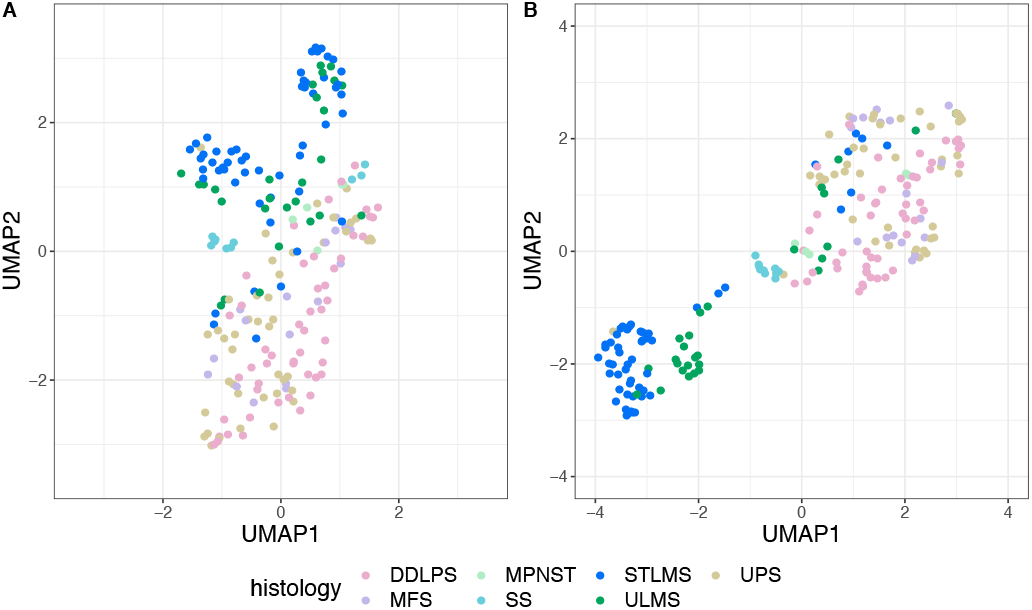
UMAP visualization of the distribution of 206 soft-tissue sarcomas, representing seven different histological subtypes (indicated with different colors) based on (A) gene targeting scores (B) expression. DDLPS: dedifferentiated liposarcoma, MFS: myxofibrosarcoma, MPNST: malignant peripheral nerve sheath tumor, SS: synovial sarcoma, STLMS: soft tissue leiomyosarcoma, ULMS: uterine leiomyosarcoma, UPS: undifferentiated pleiomorphic sarcoma.

In both the regulatory networks and expression data, the majority of leiomyosarcoma samples, represented by uterine (ULMS) and soft-tissue leiomyosarcoma (STLMS), clustered separately from other sarcoma subtypes, with a more distinct separation observed in the gene expression profiles (Figure 2). Co-localization of uterine and soft-tissue leiomyosarcomas was different between the two UMAP embeddings—while ULMS samples separated from STLMS in the expression data, clustering of leiomyosarcoma based on gene regulatory networks did not separate these subtypes. This indicates that, despite the apparent differences in gene expression between the two tissue-sites where leiomyosarcoma can develop, tumors that arise at these different sites do not have clearly distinct regulatory profiles. This analysis demonstrates that patient-specific regulatory networks capture heterogeneity among leiomyosarcoma tumors that is not directly obvious from analysis of expression data alone.

The remaining sarcoma subtypes were more spread out across the UMAP axes, with no clear co-localization of the gene regulatory networks derived from the same sarcoma histological subtype in distinct clusters, except for synovial sarcomas (SS). With the use of HDBSCAN in 2D UMAP space, we clustered the gene regulatory profiles of all 206 sarcoma samples and identified ten clusters (Supplementary Figure S1). Two of these clusters were mainly represented by leiomyosarcoma samples from mixed tissue-of-origin, confirming the heterogeneity we observed in the UMAP visualization.

### In-depth analysis of gene regulatory heterogeneity in leiomyosarcoma with PORCUPINE

The distinct regulatory clusters we identified in leiomyosarcoma motivated us to perform an in-depth analysis of the regulatory heterogeneity of leiomyosarcoma. To facilitate this, we developed a new computational tool, PORCUPINE, that can be applied to patient-specific gene regulatory networks to identify biological pathways that capture regulatory heterogeneity in a patient population (Figure 3). PORCUPINE examines regulatory co-variability of edge weights across a cohort of patient-specific networks in a pre-defined set of pathways, e.g. pathways from published resources such as Reactome [41]. The method performs PCA on all estimated regulatory interactions connected to genes from a specific pathway. It then compares the variance captured by the first principal component in the pathway to the amount of variance that would be expected by chance. This process is repeated for each pathway. Significant pathways can then be selected based on user-defined thresholds of adjusted p-value and effect size.

**Figure 3.**
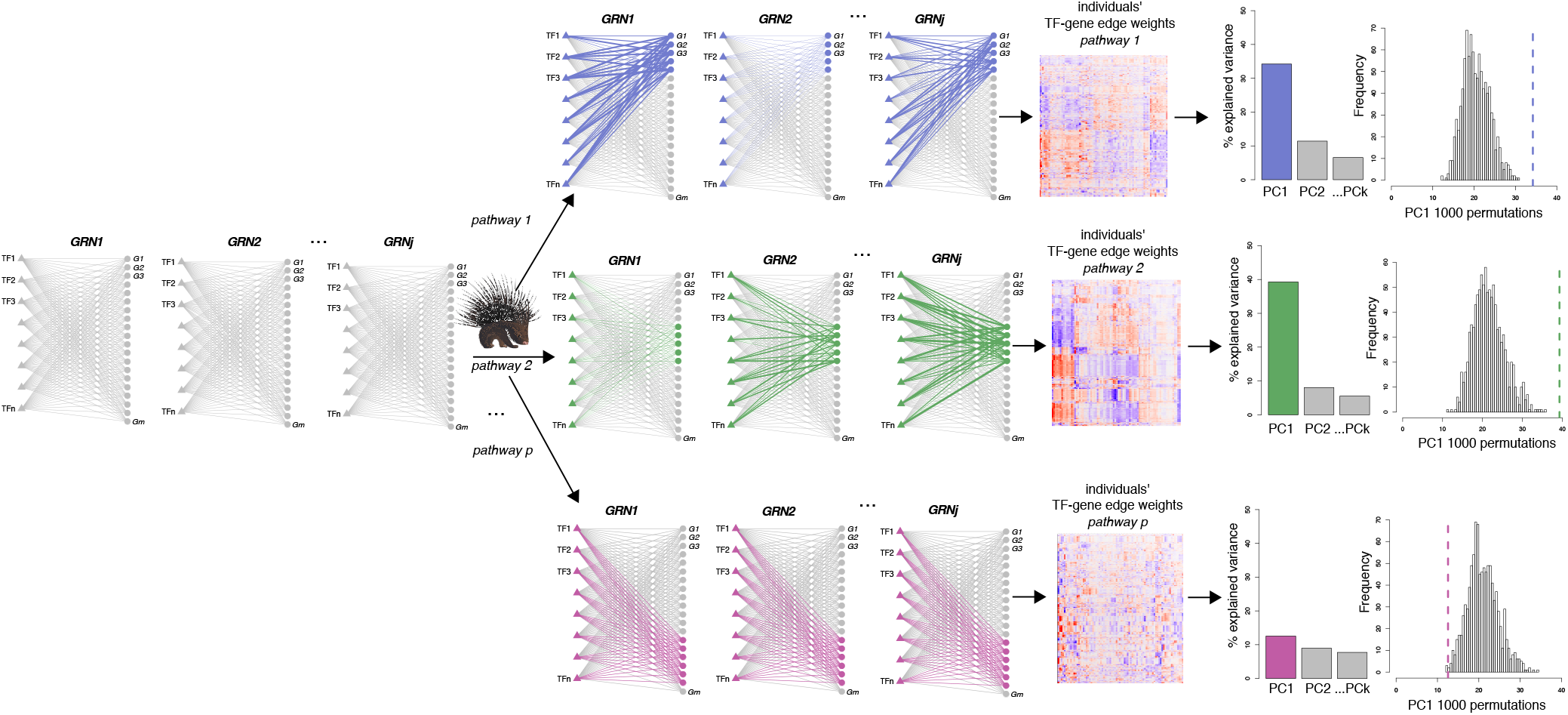
Overview of PORCUPINE (PCA to <u>O</u>btain <u>R</u>egulatory <u>C</u>ontributions <u>U</u>sing <u>P</u>athway-based Interpretation of <u>N</u>etwork <u>E</u>stimates). PORCUPINE applies the following steps: 1) TF-gene edge weight information is extracted from each individual gene regulatory network for all genes belonging to a certain pathway; 2) Principal Component Analysis is performed on the pathway-associated TF-gene weight matrix. The variance explained by the first principal component is extracted; 3) The amount of variance explained by PC1 is compared to the expected amount of variance explained, which is obtained by applying PCA on edge weights connected to 1,000 randomly generated gene sets of the same size as the selected pathway. Effect size is calculated. These steps are repeated for each pathway. P-values obtained from step 3 are then corrected for multiple testing with the Benjamini-Hochberg method.

We applied PORCUPINE to the 80 patient-specific leiomyosarcoma gene regulatory networks from TCGA, using 1,455 Reactome pathways from MSigDb (see Methods). This identified 72 significant pathways (adjusted p-value less than 0.01 and effect size >=2). We validated these results in an independent set of patient-specific networks modeled on 37 leiomyosarcoma samples from DKFZ. In the validation dataset, we identified 91 pathways, of which 37 were also identified in the networks modeled on TCGA. This overlap of 37 pathways is higher than expected by chance, with p-value <9.522e-29 based on a hypergeometric test. The pathway’s effect sizes also correlated with a Pearson correlation coefficient of 0.53. This indicates that PORCUPINE’s results are robust and highly reproducible across networks modeled on independent datasets. The 37 pathways that were detected in both datasets are visualized in Figure 4, with corresponding effect sizes.

**Figure 4.**
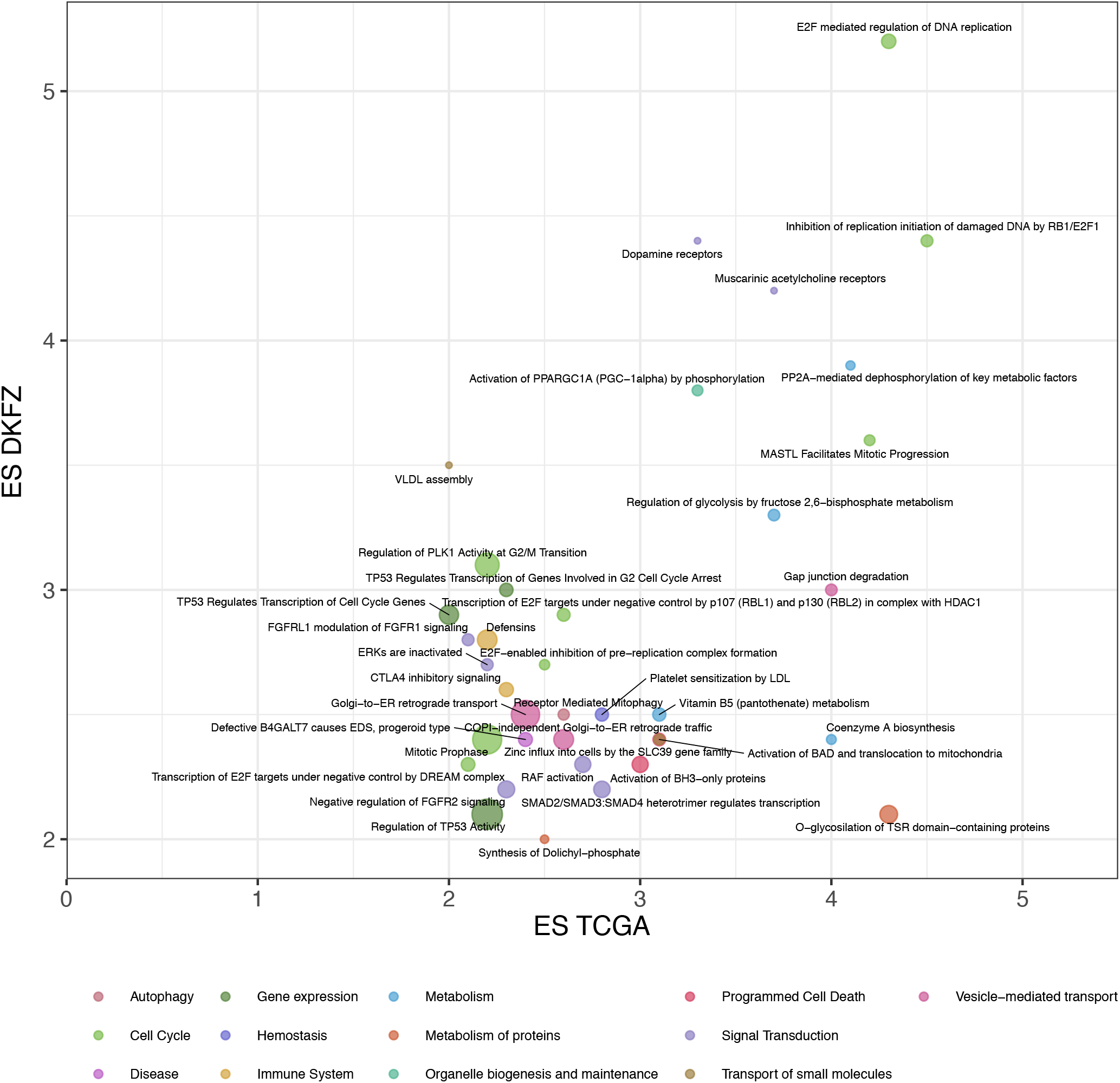
Pathways identified with PORCUPINE in both leiomyosarcoma datasets, based on FDR <0.05 and effect size >2. Pathways are colored according to their cellular function, with the size of the bubble reflecting the number of genes in the pathway.

Notably, the significant pathways varied in size, indicating that PORCUPINE analysis is not biased towards pathways of smaller or larger size (see also Supplementary Table S1).

### Regulatory heterogeneity in pathways with known and new roles in leiomyosarcoma

The two most significant pathways that were identified in both datasets are “Inhibition of replication initiation of damaged DNA by RB1/E2F1” and “E2F mediated regulation of DNA replication,” containing 13 and 22 genes, respectively. A closer examination of the genes in these pathways shows that all 13 genes in the first pathway are also part of the second pathway. PORCUPINE provides evidence of a coordinated change in the regulation of multiple genes in these pathways that is not directly captured by expression data (Supplementary Figure S2). These pathways are leiomyosarcoma-relevant, given that leiomyosarcomas are characterized by a high frequency of alterations in tumor suppressor gene *RB1*, which negatively regulates transcription factor E2F1 [18].

The 37 pathways can be further grouped into subcategories according to their cellular function (see Figure 4). Pathways with genes involved in cell cycle and signal transduction were the most frequent subcategories. Two pathways were associated with TP53 regulation, including “TP53 regulates transcription of genes involved in G2 cell cycle arrest” and “TP53 regulates transcription of cell cycle genes.” Among signal transduction pathways, we found an overrepresentation of pathways involved in Fibroblast growth factor receptors (FGFR) signaling, including “Negative regulation of FGFR2 signaling,” “FGFRL1 modulation of FGFR1 signaling,” and “ERKs are inactivated.”

FGFRs are tyrosine kinase receptors that are involved in several biological functions including regulation of cell growth, proliferation, survival, differentiation, and angiogenesis. Aberrant FGFR signaling has been shown to be associated with several human cancers and thus FGFRs are attractive druggable targets [42]. To our knowledge, among members of the FGFR family, only the inhibition of FGFR1 has been investigated in a patient with metastatic leiomyosarcoma, which showed clinical improvement [43]. There is an ongoing clinical trial testing the selective pan-FGFR inhibitor Rogaratinib to treat patients with advanced sarcoma with alterations in FGFR 1-4 [44].

Two pathways associated with immune system function were identified—“CTLA-4 inhibitory signaling” and “Defensins.” CTLA-4 is an immune checkpoint, and monocolonal antibodies such as ipilimumab and tremelimumab have been developed to target CTLA-4. These CTLA-4 inhibitors have already been used in clinical studies for treatment of several cancer types [45]. The efficacy of immunotherapy with CTLA-4 inhibitors in soft-tissue sarcoma has only been evaluated in one study to-date, in which six patients with synovial sarcoma were treated with ipilimumab [46]. To our knowledge, no clinical results testing the effect of anti-CTLA-4 in leiomyosarcoma are available or exist to-date.

To evaluate whether the identified pathways capture similar patterns of regulatory heterogeneity, we performed clustering of pathways based on inter-individual correlations of edge weights (see Methods). The pathways grouped into three main clusters, where pathways in each cluster stratified leiomyosarcoma tumors into similar subtypes. The clustering of pathways we observed was partly explained by gene overlap (Supplementary Figure S3). Pathways in cluster 1 had highest gene overlap, followed by cluster 2, with almost no gene overlap between pathways in cluster 3 (mean Jaccard indices of 0.18, 0.04, and 0.006, respectively, Supplementary Figure S3). Shared patterns of heterogeneity between pathways without apparent gene sharing can also indicate a higher order of co-regulation of these pathways. An example is the pathway “Negative regulation of FGFR2 signaling,” which belongs to cluster 1, however, based on its Jaccard indices, this pathway does not cluster with remaining pathways from the same cluster.

### Major genes and transcription factors contributing to leiomyosarcoma heterogeneity

We next identified those regulatory interactions in each of the 37 pathways that contributed most to the regulatory heterogeneity we observed in leiomyosarcoma (see Methods). Across all pathways, genes including *PPP2R1A, PPP2CB, TFDP2, CCNB1*, and *RB1* were frequently found among the top targets (Supplementary file 2). These genes are related to cell proliferation and growth. Noteworthy, *PPP2R1A* was among the top contributors in 13 out of 37 pathways and may therefore be a key player in driving leiomyosarcoma heterogeneity (see Figure 5 for its contribution to three selected pathways). It encodes for a subunit of protein phosphatase 2 (PP2), which plays a role in the negative control of cell growth and division. PP2A inactivation is a crucial step in malignant development [47]. It was previously shown that *PPP2R1A* mutation is frequent in uterine cancers [48]. However, we did not identify an association between the histological subtype of leiomyosarcoma and gene regulatory heterogeneity in pathways that had *PPP2R1A* among their major contributors. We also did not identify any significant association of patient heterogeneity scores with *PPP2R1A* mutation profiles, indicating that regulatory heterogeneity of *PPP2R1A* is not driven by somatic mutations in the gene itself.

**Figure 5.**
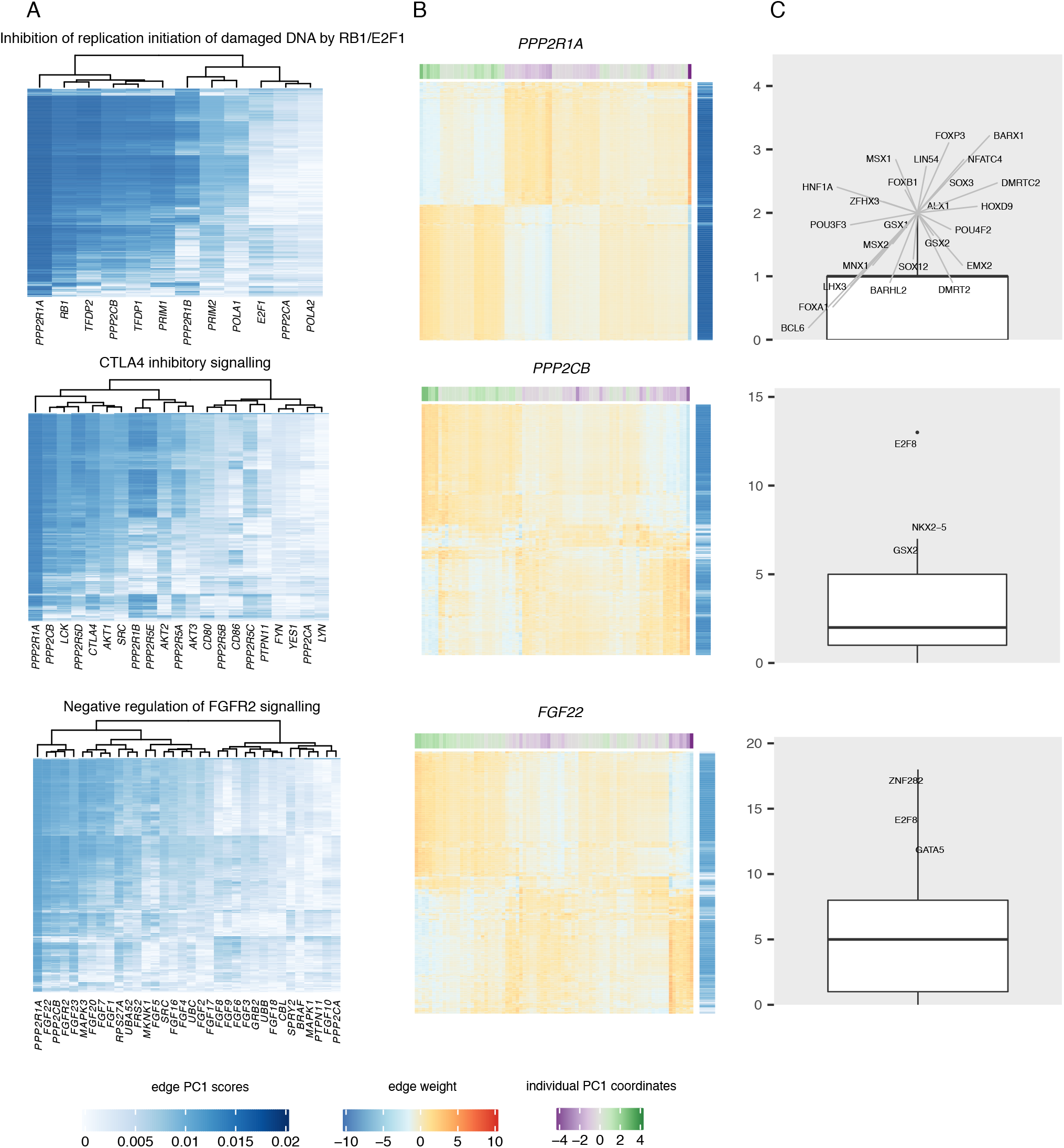
A. Heatmaps showing the contribution scores of genes and all TFs to the first principal component in three selected, significant pathways. B. Heatmaps showing the edge weights of selected genes to all TFs in these pathways. Edge weights are scaled across individuals. Row annotation shows the edge contribution scores to PC1 in each pathway. Column annotation indicates the patient heterogeneity scores in each pathway. C. Boxplots showing the number of targets for TFs with top edge contribution scores to PC1 in each pathway. TFs with a number of targets greater than the 95th percentile in each pathway are labelled.

In addition to reporting the top target genes, we identified top TFs contributing to regulatory heterogeneity in each pathway. TFs that coordinately regulated multiple targets are shown in Figure 5C for the three main pathways discussed above and in Supplementary Figure S6 for all pathways. Some TFs had a limited number of targets that they regulate in a coordinated manner, such as in the pathway “Inhibition of replication initiation of damaged DNA by RB1/E2F1,” where various TFs target a relatively low number of genes. Other TFs, such as E2F8 in “CTLA4 inhibitory signaling,” were enriched for heterogeneously targeting a large number of genes (Figure 5C, see also Supplemental Figure S4, which indicates most genes of this pathway are coordinately targeted by E2F8). E2F8 and ZNF282 were the most frequent TFs that connected to a large number of targets across many of the identified pathways (see also Supplementary Figure S6).

The E2F family of TFs contains eight members that play central roles in many biological processes, including cell proliferation, differentiation, DNA repair, cell cycle, and apoptosis. Several studies have shown that dysregulation of E2F8 is associated with oncogenesis and tumor progression in many cancers. For example, it was shown that expression of E2F8 is associated with tumor progression in breast cancer [49], human hepatocellular carcinoma [50], and lung cancer [51]. However, not much is known about the role and clinical significance of E2F8 in leiomyosaroma, nor in other sarcomas.

The role of ZNF282 (Zinc finger protein 282) in human cancers, including sarcomas, is unknown. In a study by Yeo *et al.*, it was shown that ZNF282 overexpression was associated with poor survival in esophageal squamous cell carcinoma, and depletion of ZNF282 inhibited cell cycle progression, migration, and invasion of cancer cells [52]. Additionally, the authors provided evidence that ZNF282 functions as an E2F1 co-activator, highlighting a potential connection between this TF and E2F signaling.

### Regulatory heterogeneity in leiomyosarcoma is not associated with clinical features, somatic mutations, or DNA methylation

We next explored if the heterogeneity we observed in leiomyosarcoma gene regulatory networks is associated with known features that may influence patient heterogeneity, such as clinical features and genomic data.

To investigate whether the identified pathways were associated with clinicopathological features, we performed an association analysis of the pathway-based patient heterogeneity scores with clinical features available from the TCGA and DKFZ resources (Supplementary Figure S5). There were no significant associations between the clinical features and the pathway-based patient heterogeneity scores on the first principal component (at FDR <5%). To determine whether any of the identified pathways were related to patient survival, we used the pathway-based patient heterogeneity scores on the first principal component in Cox regression models to predict patient outcome. We did not identify any significant associations with survival.

To evaluate if any of the identified pathways could classify patients with similar mutational profiles, we associated the first principal component in these pathways with pathway mutation scores. To do so, we downloaded and processed mutation data obtained from leiomyosarcoma tumors from TCGA (available for 72/80 patients) as described in Kuijjer *et al.* [32]. We performed a Kruskal Wallis test to compare the pathway-based patient heterogeneity scores on the first principal component in each of the 37 pathways between two groups, i.e. mutated compared to not mutated, for each mutated pathway. No significant differences were identified (FDR <0.05), indicating that the separation of leiomyosarcoma patients identified with PORCUPINE is independent of tumor mutation profiles. Thus, gene regulation may potentially be a new, mutation-independent mechanism driving patient heterogeneity.

To investigate if the patient heterogeneity profiles were associated with inter-individual differences in the tumor’s methylation profiles, we performed correlation analysis of the pathway-based patient heterogeneity scores on PC1 with overall DNA methylation profiles of individual tumors. There were no significant associations (FDR <0.05), indicating that regulatory heterogeneity in leiomyosarcoma is independent of methylation status.

### Regulatory heterogeneity of the identified pathways is not observed in healthy tissues

To explore if the 37 pathways we identified were cancer-specific, we assessed gene regulatory heterogeneity in healthy smooth muscle–derived tissues, represented by esophageal muscularis and uterus. In total, 283 esophageal muscularis and 90 uterus sample-specific gene regulatory networks, modeled with PANDA and LIONESS, were available from the GTEx project through the GRAND database [34]. We used PORCUPINE to characterize regulatory heterogeneity in this dataset. Among the 37 pathways identified to drive leiomyosarcoma heterogeneity, only one pathway, i.e “Gap junction degradation” was significant in these healthy tissues, indicating that 36/37 pathways we identified are leiomyosarcoma-specific and that gene regulatory heterogeneity in these pathways likely develops during sarcomagenesis.

### Regulatory heterogeneity associates with chromatin state

Finally, we investigated whether network heterogeneity corresponds to chromatin accessibility. To do so, we profiled RNA-seq and ATAC-seq for four leiomyosarcoma cell lines. To translate our findings on the inter-patient heterogeneity in leiomysarcoma to the cell lines, we constructed cell line specific gene regulatory networks based on the RNA-seq data, and placed these networks on the regulatory map of leiomyosarcoma patients. Cell lines clustered among the DKFZ-LMS patient specific networks (Supplementary figure S8), and we could confirm 29/37 pathways when we included these cell lines in our analyses.

We next clustered ATAC-seq profiles of the four cell lines (three replicates for each cell line, see Supplementary Figure S7). The cell lines had distinct chromatin profiles with SK-LMS-1 and MES-SA clustering separately from SK-UT-1 and SK-UT1-B—two cell lines that are derived from the same donor. We then assessed whether promoters of genes from the significant pathways detected by PORCUPINE are located within open chromatin regions. To do so, we compared peak accessibility at the promoters of these genes to that at promoters of randomly selected genes. We observed a significant enrichment in open chromatin regions for the heterogeneously regulated genes (Figure 6A). In addition, we compared expression of these genes to randomly selected gene sets, and found they are also highly expressed (Figure 6B). This suggests that genes that are located in open chromatin regions are more likely to be regulated by different sets of TFs, which could have implications for network-based biomarker detection or the development of subtype-specific targets for treatment.

**Figure 6.**
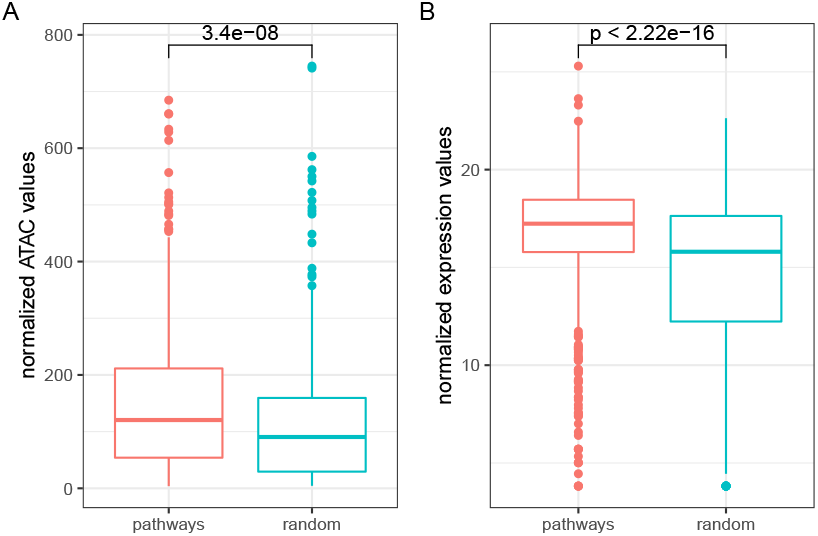
Comparison of chromatin accessibility of promoter regions of the genes and expression of the genes in pathways identified by PORCUPINE across four cell lines to random genes.

Finally, we evaluated whether genes from the top heterogeneously regulated pathway “E2F mediated regulation of DNA replication” were also over-represented in differentially accessible regions between leiomyosarcoma cell lines. To do so, we called differentially accessible regions in pairwise comparisons between the four cell lines (six comparisons in total). The promoter of *PPP2R1A*, the gene we found to be most enriched for heterogeneous regulation in the two patient cohorts, was differentially accessible in all pairwise comparisons between cell lines, except between SK-UT-1 and SK-UT1-B (see Supplementary Table S2). However, as these two cell lines are derived from the same donor, they are expected to have comparable regulatory profiles. This indicates that the differential heterogeneity observed in the patient-specific regulatory networks are not only associated with open chromatin states in general, but also with more subtle differences in chromatin landscapes between individual tumors.

## Discussion

In this work, we hypothesized that classification of soft-tissue sarcoma patients on the basis of gene regulatory networks has the potential to provide additional, novel information to stratify patients into clinically meaningful subgroups, to point to potential new targets for treatment, and to identify new biomarkers to guide selecting patients most likely to benefit from a specific treatment.

To this end, we developed PORCUPINE, a novel computational approach to map heterogeneity of gene regulation across a patient population. We applied the method to model heterogeneity of gene regulation in leiomyosarcoma, which we found to present a high level of heterogeneity in a pan-sarcoma network analysis. Applying PORCUPINE to two independent leiomyosarcoma cohorts identified 37 pathways that robustly capture gene regulatory heterogeneity in the disease. Among the detected pathways, we identified pathways that could represent potential targets for treatment of subgroups of leiomyosarcoma patients, including RB1/E2F1 signaling, pathways involved in FGFR signaling, and CTLA4 inhibitory signaling. While these pathways have been described as potential targets for treament of sarcomas, not all patients may respond to such approaches, as, for example, was recently shown for treatment with a CTLA4 inhibitor in synovial sarcoma [46]. Stratifying patients based on the regulatory profiles of these pathways could potentially help identify subgroups of patients that are likely to respond to treatments that act on these pathways.

PORCUPINE highlighted genes and TFs that are enriched in driving heterogeneity among leiomyosarcoma patients, including *RB1* and *PPP2R1A* as target genes, as well as the TFs E2F8 and ZNF282, which could potentially be inhibited [53]. Through gene regulatory network modeling and ATAC-seq profiling in leiomyosarcoma cell lines, we found that promoters of the most heterogeneously regulated genes in leiomyosarcoma are enriched for open chromatin states. This suggests that genes in open chromatin states may be more prone to receive differential binding by TFs, which could have implications for the detection of regulatory biomarkers or subtype-specific targets for treatment.

We performed our study on four leiomyosarcoma cell lines that are also represented in extensive cell profiling and functional genomics initiatives such as DepMap from Broad Institute. While we could capture heterogeneous regulation in most of the identified pathways, the small number of cell lines may likely not fully represent the landscape of heterogeneous gene regulation we observed in the patient cohorts, which is a limitation of our study. However, we could still identify significant differential chromatin states for the top heterogeneously regulated gene in the patient population, *PPP2R1A*, indicating that our network models may potentially also capture subtle differences in chromatin states in a patient population.

We developed PORCUPINE as user-friendly R package that can be applied to single-sample networks. While similar approaches have previously been successfully applied to study heterogeneity in cancer using gene expression profiles [54], our approach differs from these methods as we specifically designed it to analyze large-scale, genome-wide gene regulatory networks. Of note, while we used PORCUPINE on networks modeled with PANDA and LIONESS, the tool is not limited to these specific methodologies, and could potentially also be used to analyze (bipartite) networks modeled with other single-sample approaches. Of course, when applying PORCUPINE, one should consider cohort sample size as well as the use of an independent validation dataset, as we showed here by including an independent leiomyosarcoma dataset, which are both important to include to detect relevant and robust pathways. Additionally, it is important to note that, while the use of a large set of randomized pathways is beneficial, it comes with disadvantage of an increase in computational load.

Genome-wide gene regulatory networks represent high-dimensional data. Usually, network summary statistics, such as gene targeting scores, closeness centrality, or betweenness centrality, are calculated prior to any further analysis to reduce the dimensions of large-scale networks. Then, to identify heterogeneity across a cohort, unsupervised clustering approaches are widely used [55]. The advantage of PORCUPINE is that it can be directly applied to high-dimensional networks, as it uses as input the network’s edge weights instead of a summary statistic. Moreover, as it does this per individual biological pathway, the output is not just a collection of significant differential edges that need to be further analyzed, but rather a list of differentially regulated pathways that are easy to interpret. Additionally, the method can capture significant aspects of heterogeneity among individuals in situations when no clear population structure with well defined clusters can be revealed. PORCUPINE estimates pathway-based patient heterogeneity scores that can facilitate the identification of either continuous gradients or discrete gene regulatory subtypes and that can be further used in association analyses with clinical covariates, or in survival analyses, as we have shown in this work.

In summary, with PORCUPINE, we uncovered patterns of inter-patient heterogeneity at the level of transcriptional regulation in tumors and cell models, and identified genes and pathways that may represent therapeutic entry points in leiomyosarcoma. Our approach thereby provides one of the first steps towards implementing network-informed personalized medicine in soft-tissue sarcomas.

## ACKNOWLEDGMENTS

This work was supported by the Norwegian Research Council, Helse Sør-Øst, and University of Oslo through the Centre for Molecular Medicine Norway (187615, to MLK), the Norwegian Research Council (313932, to MLK), Familien Blix Fond (to TB and MLK), as well as the Emmy Noether Programme Grant from the German Research Foundation (DFG, No. CH 2302/1-1, to PC). The authors would like to thank Jing Yang and the Omics IT and Data Management Core Facility (ODCF) of the DKFZ for help with data preprocessing and transferring, Romana Pop for testing the PORCUPINE code, and Ingrid Kjelsvik and Elisa Bjørgo for administrative support.

## SUPPLEMENTARY FIGURES AND TABLES

**Supplemental Table S1.** Proportions of pathways of different sizes among Reactome pathways and pathways identified with PORCUPINE in the TCGA-LMS and DKFZ-LMS datasets.

| ptw genes | Reactome% | TCGA% | DKFZ% |
| --- | --- | --- | --- |
| <50 | 75.8 | 81.9 | 78 |
| $\geq 50 \& \leq 100$ | 16 | 12.5 | 12 |
| >100 & <150 | 5.9 | 4.2 | 5.8 |
| $\geq 150$ | 2.3 | 1.4 | 4.4 |

**Supplemental Table S2.** Genes from the “E2F mediated regulation of DNA replication” pathway mapped to differential accessible regions (DARs) in each of the pairwise cell line comparisons.

| Comparison | Among DARs | Not among DARs |
| --- | --- | --- |
| SK-UT-1B vs SK-LMS-1 | *PPP2R1B, PPP2R1A, E2F1, PRIM2, MCM8, TFDP2, POLA2* | *PPP2CA, PRIM1, CCNB1, ORC4, POLA1, ORC1, PPP2CB, CDK1, ORC5, RB1, TFDP1, ORC2* |
| SK-UT-1B vs MES-SA | *PPP2R1A, ORC5, ORC1, CCNB1, ORC2, ORC4, RB1, PPP2R1B* | *POLA1, TFDP1, PRIM2, PPP2CA, PRIM1, TFDP2, POLA2, MCM8, PPP2CB, E2F1, CDK1* |
| SK-UT-1 vs SK-UT-1B | *CDK1, PRIM1, ORC4* | *CCNB1, PPP2CA, ORC2, TFDP2, PPP2R1A, ORC1, TFDP1, PPP2R1B, POLA1, E2F1, RB1, ORC5, PPP2CB, PRIM2, POLA2, MCM8* |
| SK-UT-1 vs SK-LMS-1 | *PPP2R1A, PPP2R1B, E2F1, PRIM2, MCM8, PRIM1, PPP2CA, CCNB1, CDK1, POLA2, TFDP2, POLA1* | *TFDP1, ORC4, PPP2CB, ORC5, ORC2, RB1, ORC1* |
| SK-UT-1 vs MES-SA | *CCNB1, PPP2R1A, ORC4, ORC1, ORC2, ORC5, PRIM1, CDK1, PPP2R1B, TFDP1, POLA1, RB1, PRIM2* | *POLA2, PPP2CA, PPP2CB, MCM8, E2F1, TFDP2* |
| MES-SA vs SK-LMS-1 | *PPP2R1B, PRIM2, E2F1, ORC5, MCM8, ORC1, ORC4, ORC2, POLA2, PPP2CA, PPP2R1A, RB1, TFDP2, CCNB1* | *TFDP1, POLA1, CDK1, PRIM1, PPP2CB* |

**Supplementary Figure S1.**
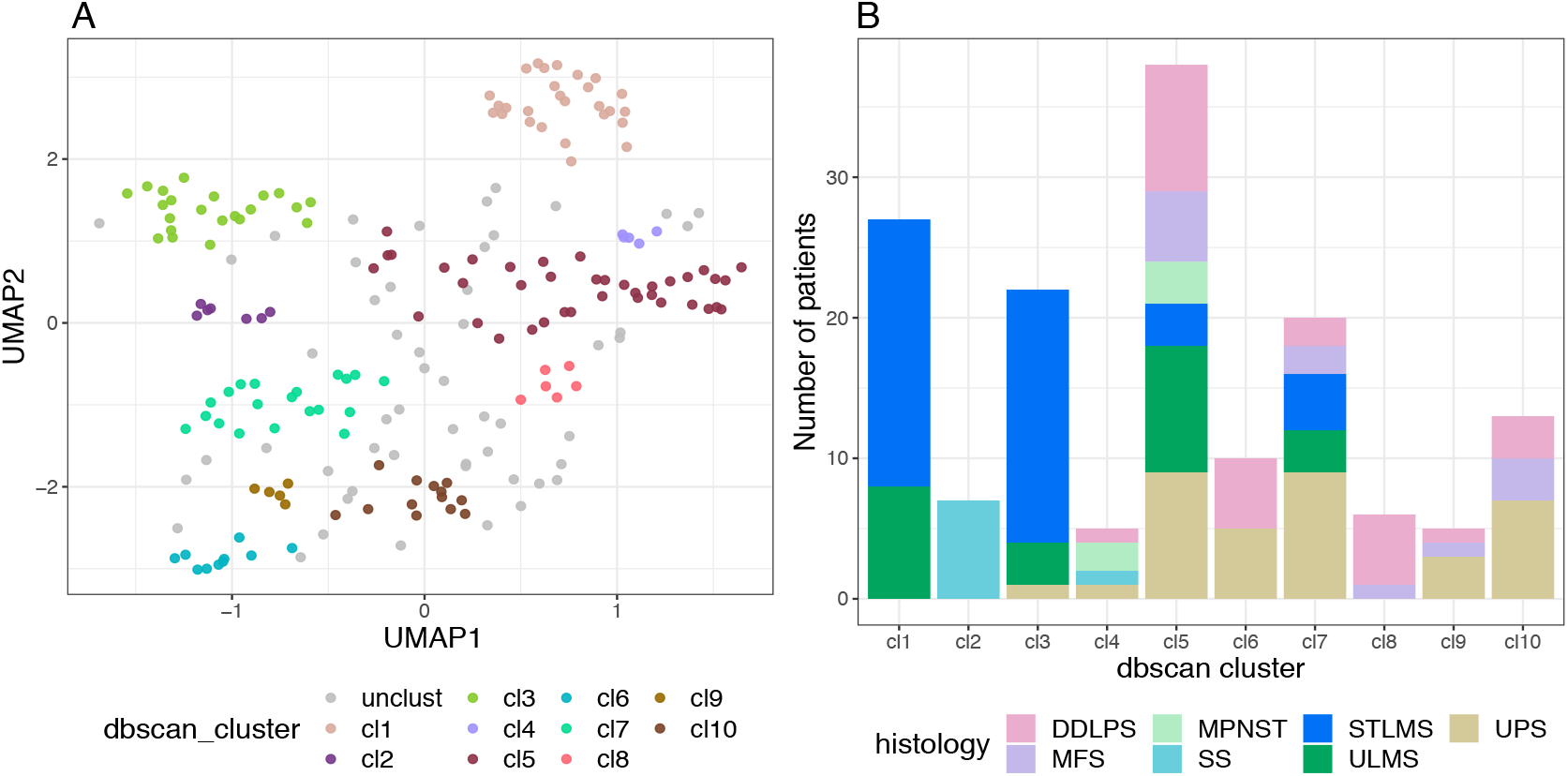
A. HDBSCAN clustering of 206 soft-tissue sarcomas on the first two UMAP dimensions obtained from applying UMAP on gene targeting scores. B. Distribution of STS histological subtypes across HDBSCAN clusters. unclust: samples that did not cluster with any subtype, cl: cluster. DDLPS: differentiated liposarcoma, MFS: myxofibrosarcoma, MPNST: malignant peripheral nerve sheath tumor, SS: synovial sarcoma, STLMS: soft tissue leiomyosarcoma, ULMS: uterine leiomyosarcoma, UPS: undifferentiated pleiomorphic sarcoma.

**Supplementary Figure S2.**
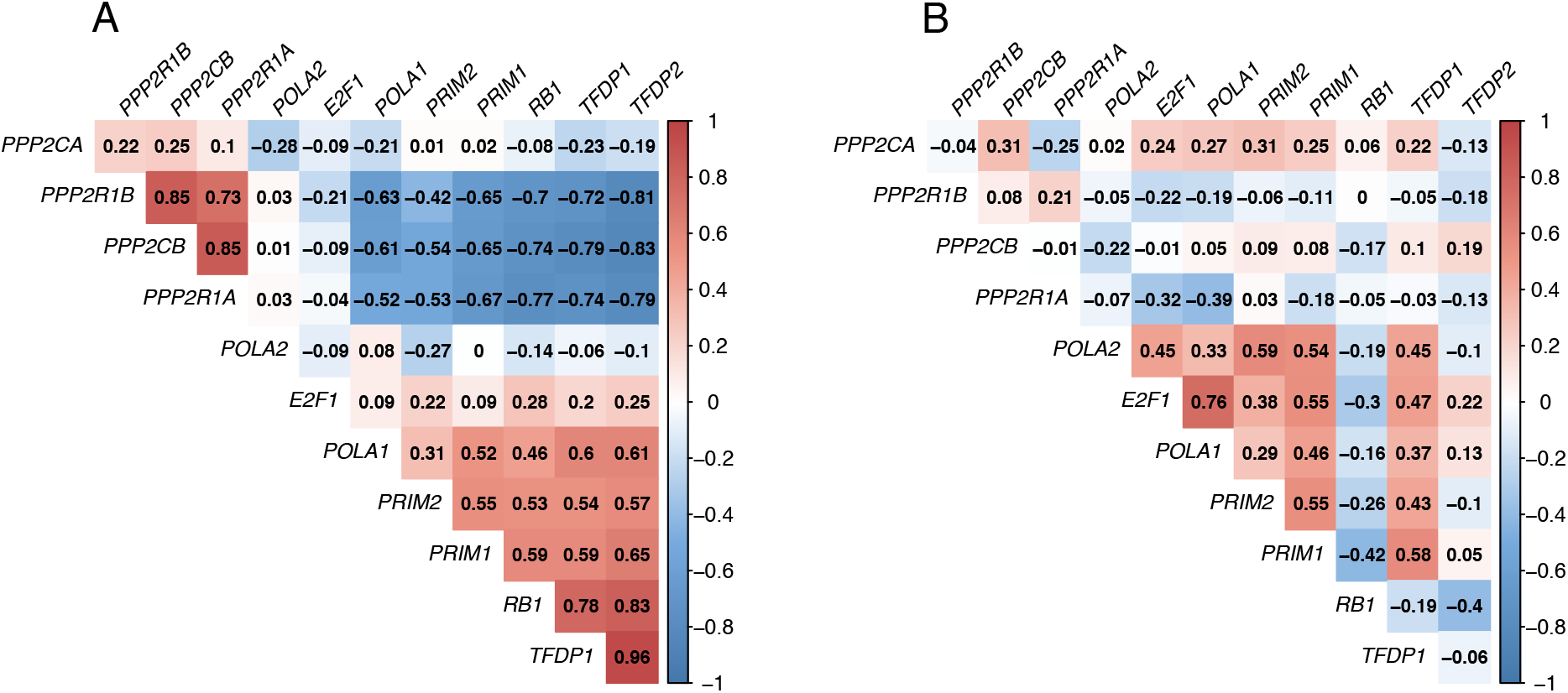
Pearson correlations between A. targeting scores and B. expression levels of genes belonging to the pathway “Inhibition of replication initiation of damaged DNA by RB1/E2F1.”

**Supplementary Figure S3.**
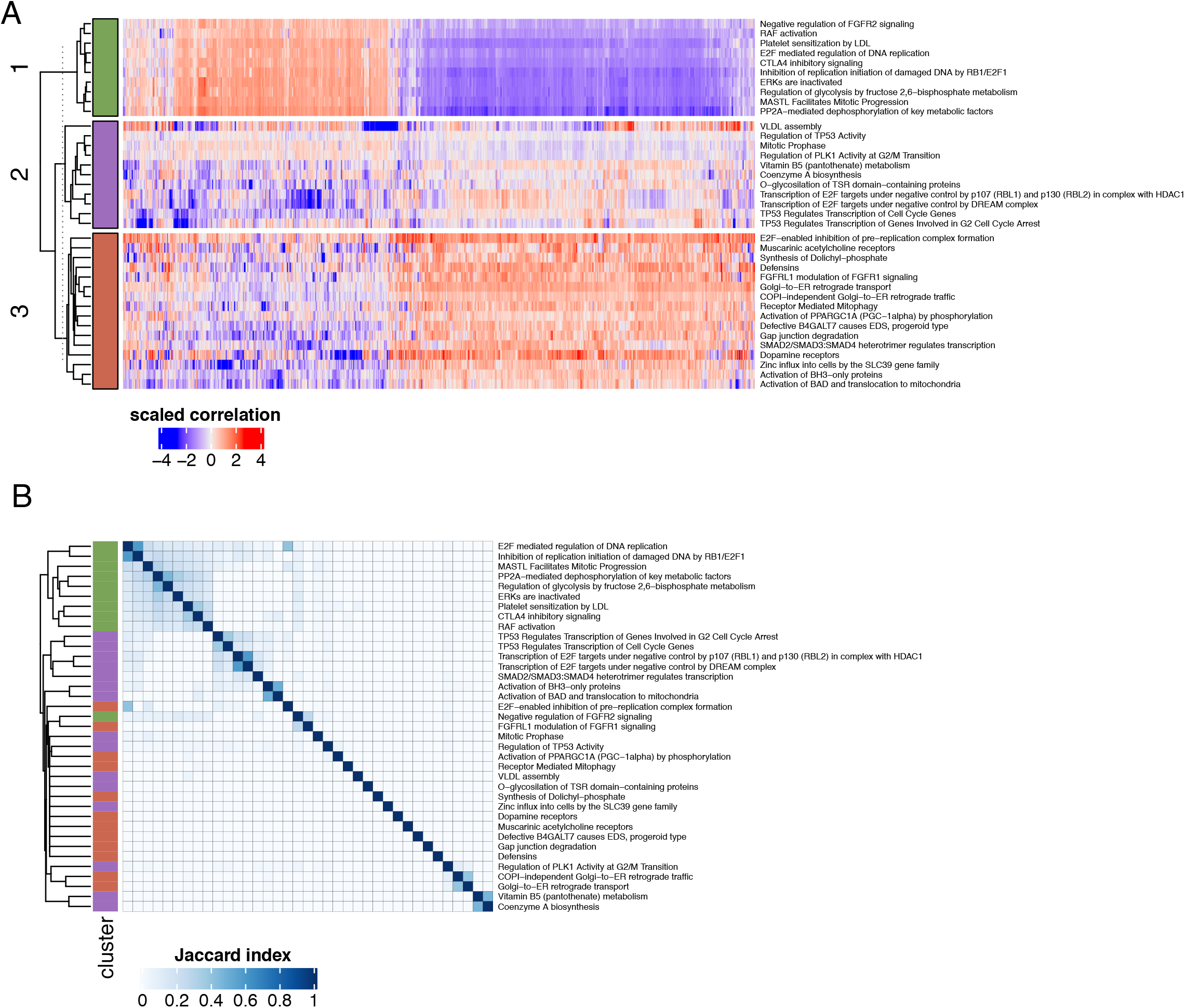
Clustering of 37 pathways based on A. pair-wise correlations between individual networks from the TCGA-LMS dataset; and B. the proportion of shared genes between the pathways. The “scaled correlation” in A indicates z-scored normalized Pearson correlations. The color code of clusters in Figure B corresponds to the color code used in Figure A.

**Supplementary Figure S4.**
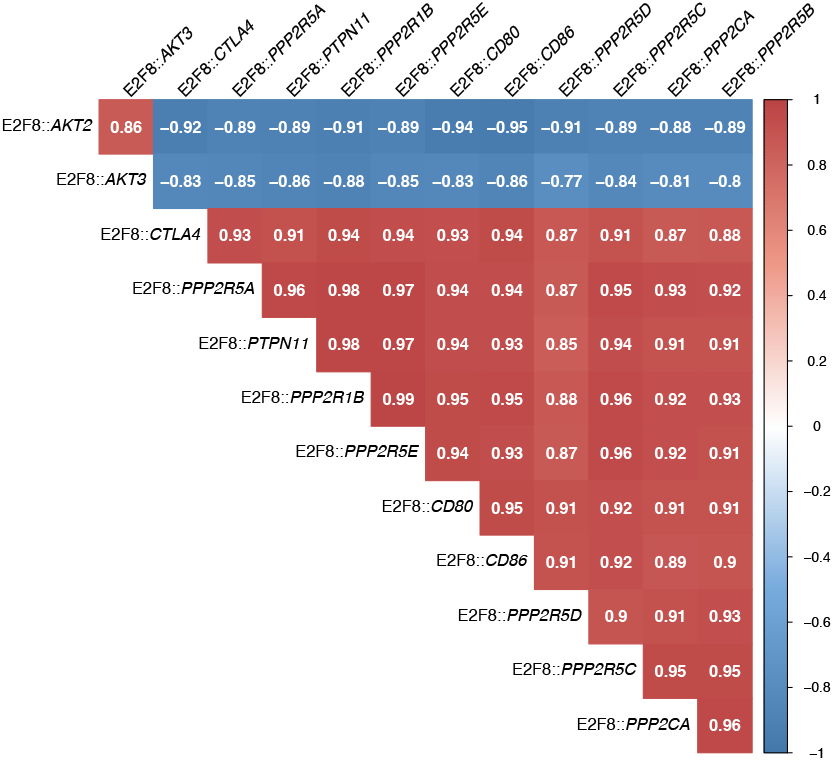
Pearson correlations among edge weights of target genes of the transcription factor “E2F8” in the pathway “CTLA4 inhibitory signalling.”

**Supplementary Figure S5.**
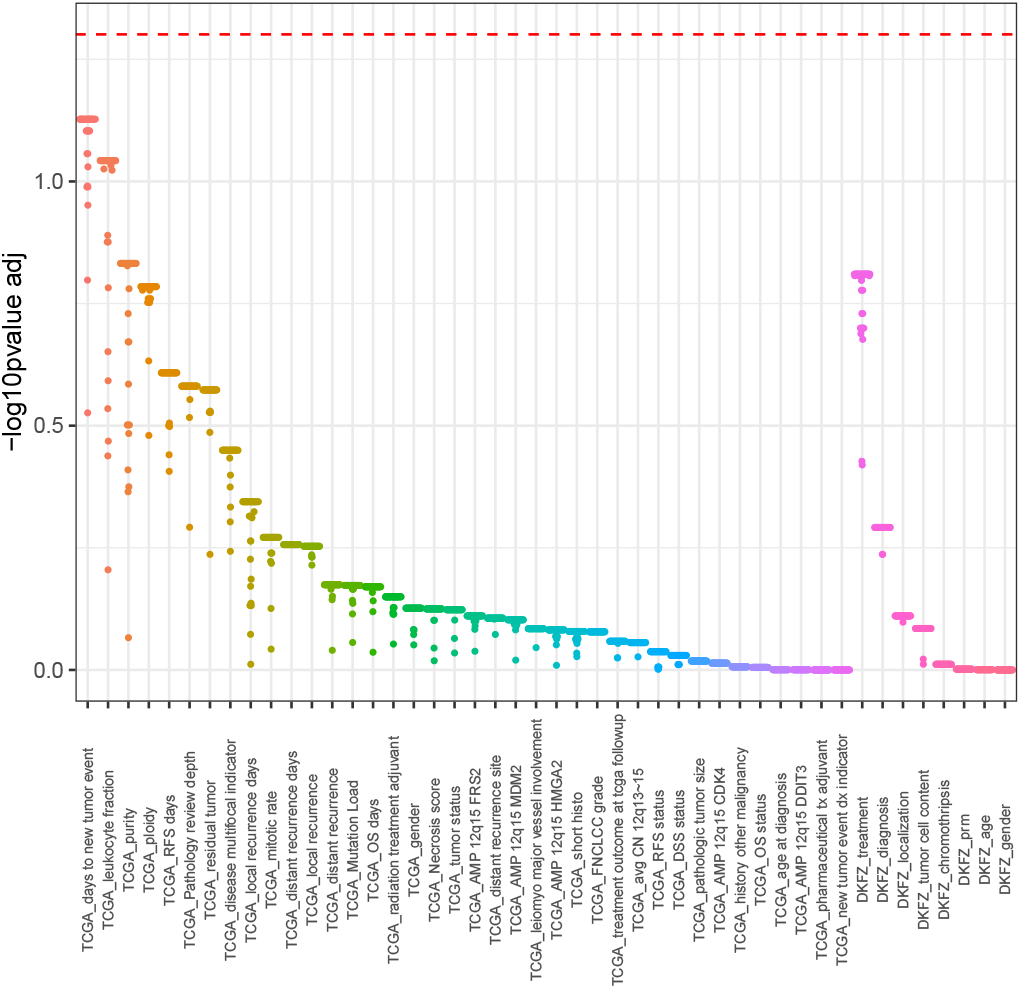
Association of the clinical features of patients and pathway-based patient heterogeneity scores on PC1 in each of the 37 pathways. Associations are shown for both cohorts (TCGA and DKFZ). The y-axis indicated the negative log base 10 of the FDR-adjusted p-value. The dotted line indicates a p-value of 0.05.

**Supplementary Figure S6.**
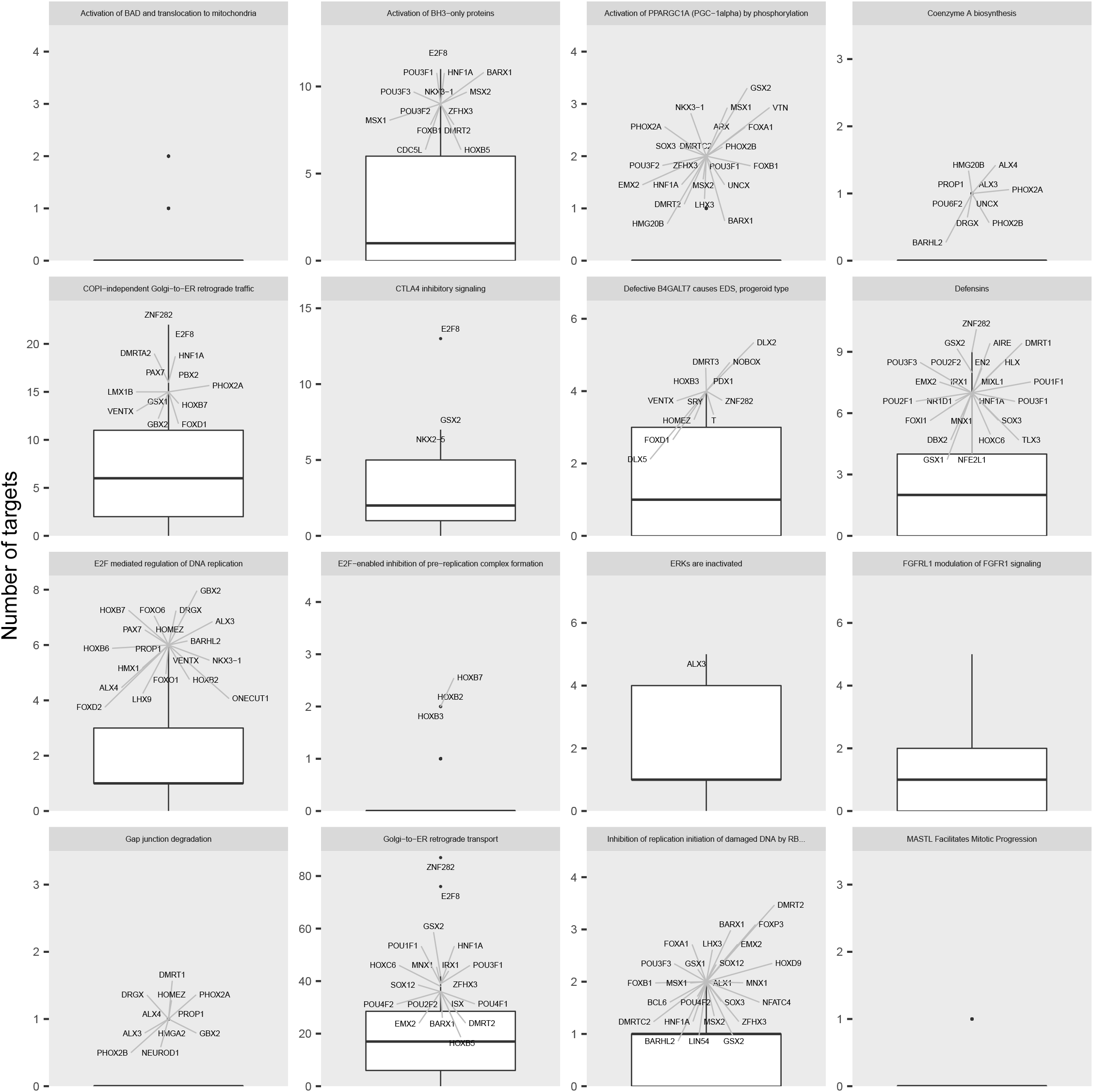
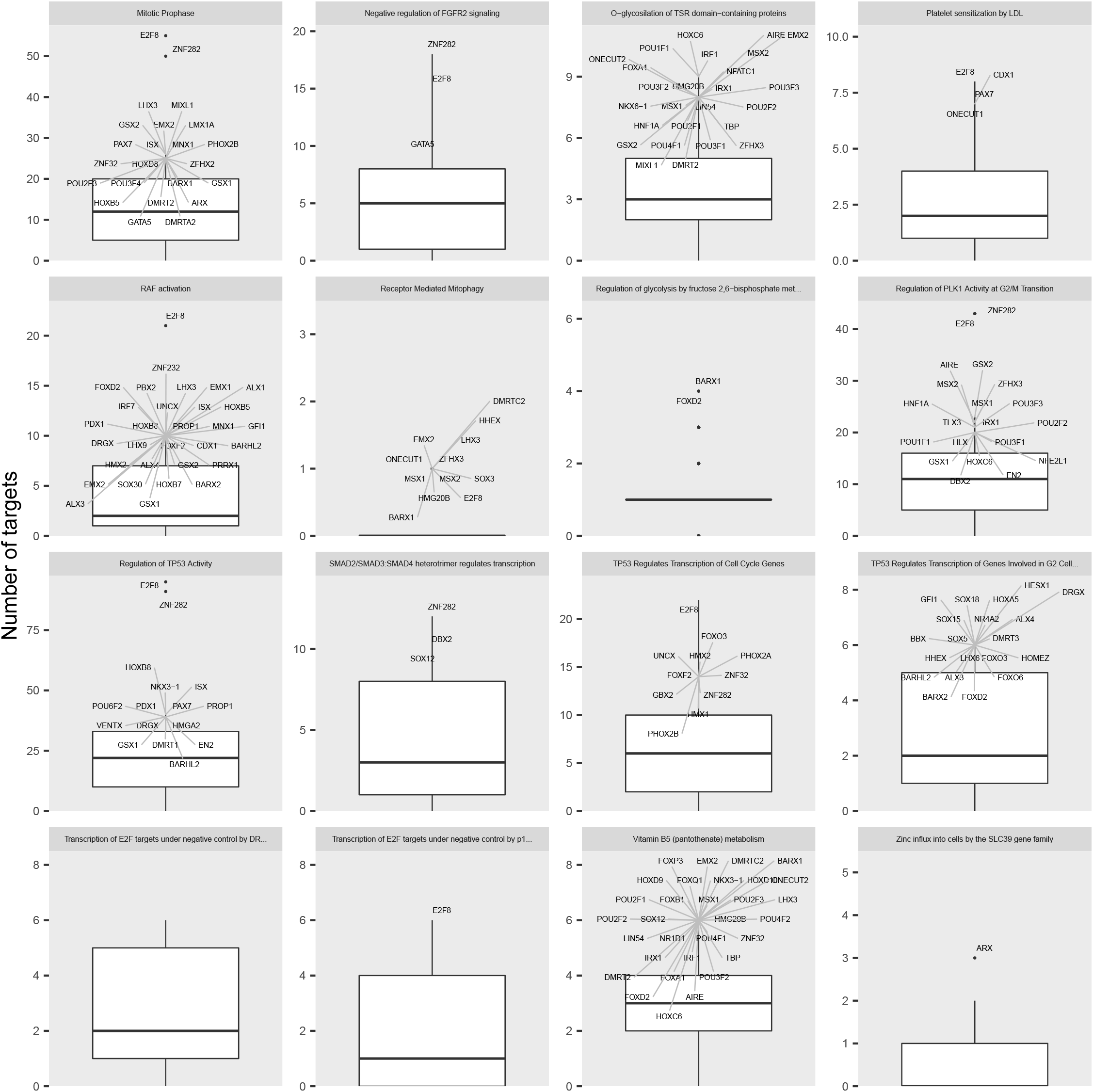
Boxplots showing the number of targets for TFs with most highly weighted values to PC1 in each pathway. 32 out of 37 pathways had edge weights with contribution scores above the threshold. TFs with a number of targets greater than the 95th percentile are labelled.

**Supplementary Figure S7.**
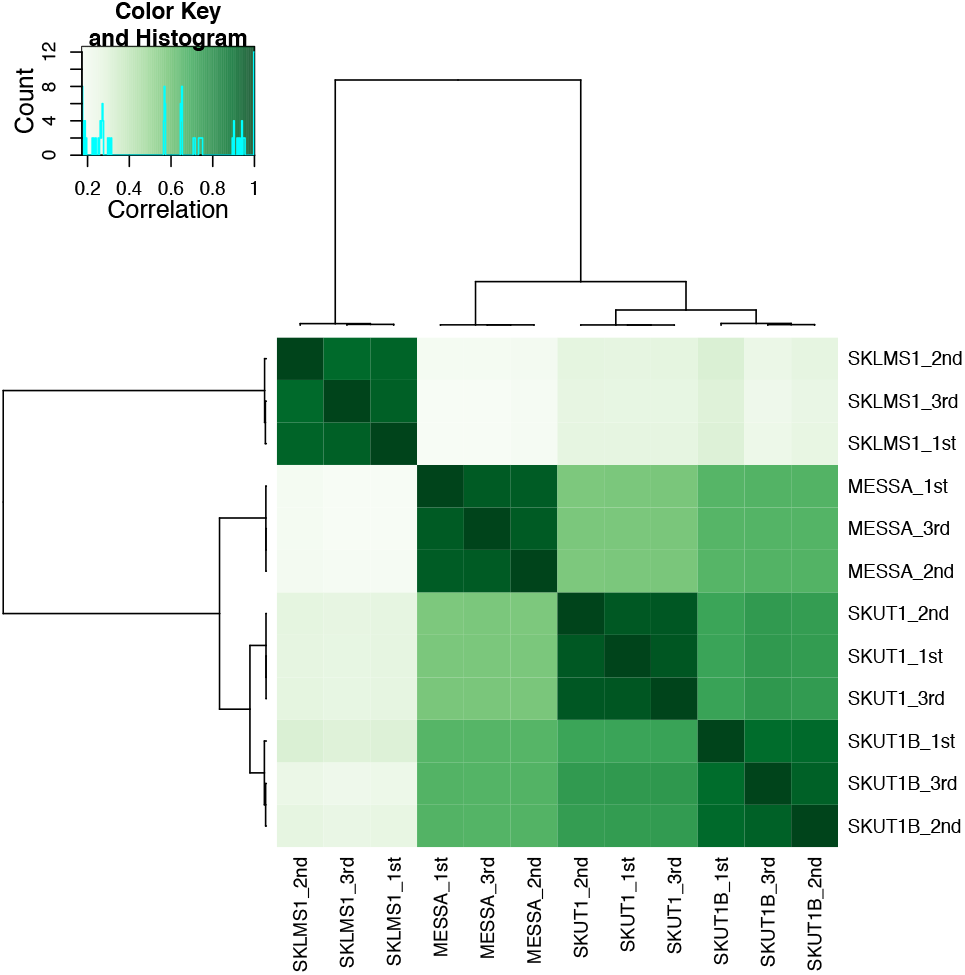
Hierarchical clustering of cell lines based on their ATAC-seq profiles. Numbers indicate technical replicates. Correlation (color key) indicates the Pearson correlation coefficient.

**Supplementary Figure S8.**
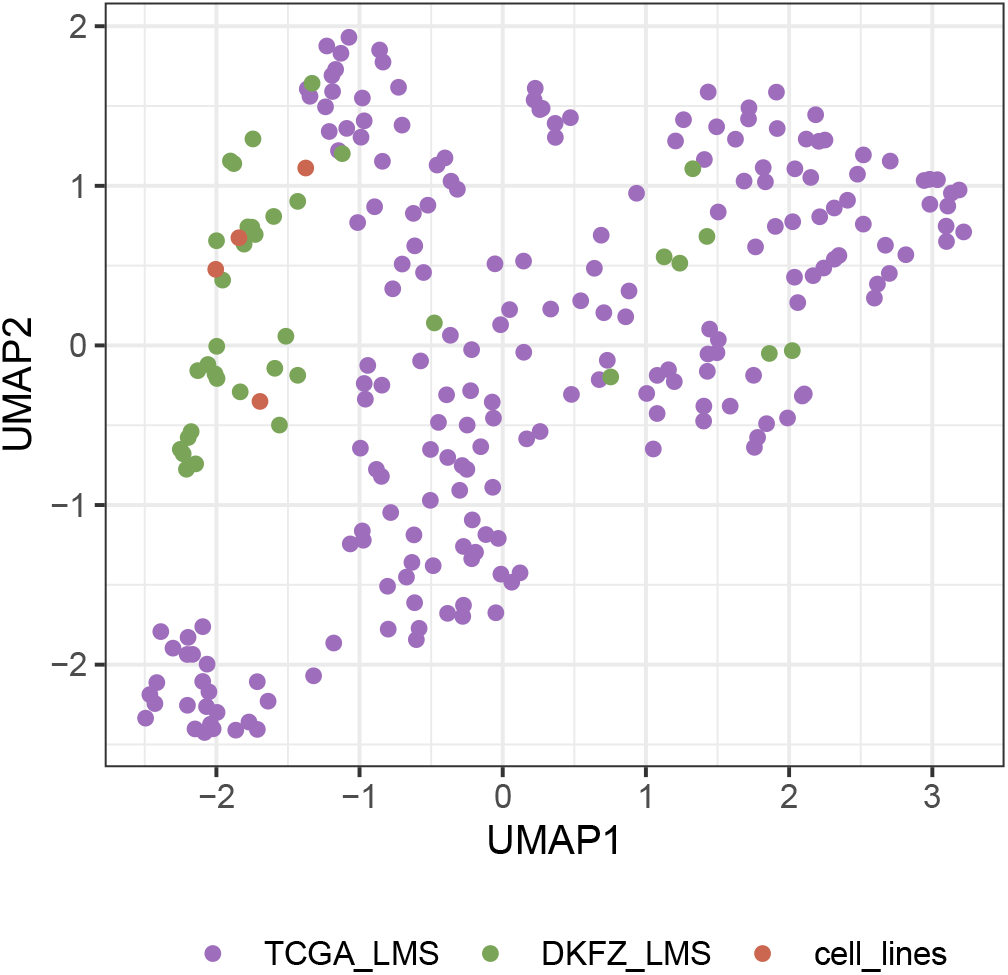
UMAP visualization of the distribution of leiomyosarcomas from three different datasets (indicated with different colors).

