## Supplementary File 1 for "Heterogeneity in the gene regulatory landscape of leiomyosarcoma"

**OMNI-ATAC protocol**

**Reagents, buffers, solutions, and DNA oligos**

A. **Reagents**

- Acetic acid (Sigma-Aldrich, #695092-100ML)
- AMPure XP (Beckman Coulter, #A63880)
- Buffer EB (Qiagen, #19086)
- Digitonin (Promega, #G9441)
- Dimethylformamide (Merck Millipore, #103034)
- DMSO (Sigma-Aldrich, #D8418)
- 0.5 M EDTA solution pH 8.0 (Enzo Life Sciences, #JBS-BU-105)
- Ethanol (Carl Roth, #T171.3)
- Guanidine thiocyanate (Sigma-Aldrich, #G9277)
- Hard-Shell 96-Well PCR Plates (Bio-Rad, #HSP9601)
- High Sensitivity D1000 ScreenTape (Agilent Technologies, #5067-5584)
- High Sensitivity D1000 Reagents (Agilent Technologies, #5067- 5585)
- Illumina NextSeq 550 Paired-End 75 bp Mid-Output (104 M reads)
- ILMN Tag DNA Enzyme & Buffer Small Kit (Illumina, #20034197)
- Loading Tips (1EA/PK) (Agilent Technologies, #5067-5153)
- MgCl2 (Sigma-Aldrich, #63069-100ML)
- Microseal 'B' PCR Plate Sealing Film, adhesive, optical (Bio-Rad, #MSB1001)
- Mx3000P Strip Tubes (Agilent Technologies, #401428)
- Mx3000P Optical Strip Caps (Agilent Technologies, #401425)
- NEBNext High-Fidelity 2x PCR Master Mix (New England Labs, #M0541S)
- NP-40 (10% in H₂O) (BioVision, #BV-2111-100)
- Polyethylene glycol 8000 (Carl Roth, #0263.1)
- PBS buffer (Gibco, #10010023)
- Sequencing primers (Sigma-Aldrich)
- Qubit dsDNA HS assay kit (Invitrogen, #Q32851)
- Qubit Assay Tubes-500 tubes (Invitrogen, #Q32856)
- Sodium Chloride Stock Solution (5M) (Biomol, #Cay600211-100)
- SYBR Green I (Invitrogen, # S-7563)
- Tris (Bio-Rad, #1610716)
- Tris-HCl Buffer 1M pH 7.4 (Biotrend, #21420063-1)
- Tween 20 (Sigma-Aldrich, #P9416)
- UltraPure DNase/RNase-Free Distilled Water-500 mL (Invitrogen, #10977035)

B. **Buffers**

| **ATAC-Resuspension Buffer (RSB)** | | |
| --- | --- | --- |
| **Component** | **Final conc.** | **For 10 mL** |
| 1 M Tris-HCl pH 7.4 | 10 mM | 100 μL |
| 5 M NaCl | 10 mM | 20 μL |
| 1 M MgCl_2_ | 3 mM | 30 μL |
| Sterile Milli Q water | N/A | Up to 10 mL |

**Note:** filter sterilize (32 mm filter) and store at 4°C

| **10X TMgAc Buffer** | | |
| --- | --- | --- |
| **Component** | **Final conc.** | **For 10 mL** |
| 1 M Tris acetate pH 7.6 | 100 mM | 1 mL |
| 1 M MgCl_2_ | 50 mM | 500 μL |
| Sterile Milli Q water | N/A | Up to 10 mL |

**Note:** filter sterilize (32 mm filter) and store at 4°C

| **5X TMgAc-DMF Buffer** | | |
| --- | --- | --- |
| **Component** | **Final conc.** | **For 2 mL** |
| 10X TMgAc Buffer | 5X | 1 mL |
| Dimethylformamide | 50% v/v | 1 mL |

**Note:** prepare in a sterile glass bottle

| **AMPure buffer** | | |
| --- | --- | --- |
| **Component** | **Final conc.** | **For 50 mL** |
| PEG 8000 | 18% w/v | 9 g |
| 5M NaCl | 2.5 M | 25 mL |
| 1M Tris-HCl pH 8.0 | 10 mM | 500 µL |
| 0.5M EDTA | 1 mM | 100 µL |
| Tween-20 | 0.05% | 25 µL |
| Sterile Milli Q water | N/A | Up to 50 mL |

**Note:** filter sterilize (0.20 µm filter), treat with UV light for 1 h, and store at 4°C in aliquots (5 mL

each) up to 1 month

| **Lysis buffer A** | |
| --- | --- |
| **Component** | **x1** |
| ATAC-RSB | 48.5 μL |
| 10% NP-40 | 0.5 μL |
| 10% Tween 20 | 0.5 μL |
| 1% digitonin | 0.5 μL |
| **Total** | **50 μL** |

| **Lysis buffer B** | |
| --- | --- |
| **Component** | **x1** |
| ATAC-RSB | 495 μL |
| 10% Tween 20 | 5 μL |
| **Total** | **500 μL** |

| **Transposition buffer** | |
| --- | --- |
| **Component** | **x1** |
| 5X TMgAC-DMF | 10 μL |
| 1X PBS | 16.5 μL |
| 10% Tween 20 | 0.5 μL |
| 1% digitonin | 0.5 μL |
| Nuclease-free water | 20 μL |
| **Total** | **47.5 μL** |

C. **Solutions**

- Digitonin: dilute 1:1 in nuclease-free water (1%); store at -20 °C in aliquots (5 μL each) up to 6 months and avoid more than 5 freeze-thaw cycles
- Tween 20: prepare a 10% v/v Tween 20 solution in sterile Milli Q water; store at 4 °C in aliquots (500 μL each)
- NP-40: prepare a 10% v/v NP-40 solution in sterile Milli Q water; store at 4 °C in aliquots (500 μL each)
- Guanidine thiocyanate: prepare a 5 M solution in sterile Milli Q water and filter sterilize it (32 mm filter); store at -20 °C in aliquots (500 μL each) and avoid more than 5 freeze-thaw cycles

D. **DNA oligos**

| **Primer** | **Sequence** |
| --- | --- |
| Tn5mCP1n | **AATGATACGGCGACCACCGAGATCTACAC**TCGTCGGCAGCGTC |
| Tn5mCBar6 | CAAGCAGAAGACGGCATACGAGAT**CATGTCTCA**GTCTCGTGGGCTCGG |
| Tn5mCBar8 | CAAGCAGAAGACGGCATACGAGAT**GTATCAGTC**GTCTCGTGGGCTCGG |
| Tn5mCBar9 | CAAGCAGAAGACGGCATACGAGAT**TCGCCTTA**GTCTCGTGGGCTCGG |
| Tn5mCBar10 | CAAGCAGAAGACGGCATACGAGAT**CTAGTACG**GTCTCGTGGGCTCGG |
| Tn5mCBar12 | CAAGCAGAAGACGGCATACGAGAT**GCTCAGGA**GTCTCGTGGGCTCGG |
| Tn5mCBar16 | CAAGCAGAAGACGGCATACGAGAT**CCTCTCTG**GTCTCGTGGGCTCGG |
| Tn5mCBar19 | CAAGCAGAAGACGGCATACGAGAT**TGCCTCTT**GTCTCGTGGGCTCGG |
| Tn5mCBar20 | CAAGCAGAAGACGGCATACGAGAT**TCCTCTAC**GTCTCGTGGGCTCGG |

Purification method: HPLC (resuspended in 1X TE buffer to a final concentration of 100 μM)
