## Supplementary File 2 for "Heterogeneity in the gene regulatory landscape of leiomyosarcoma"

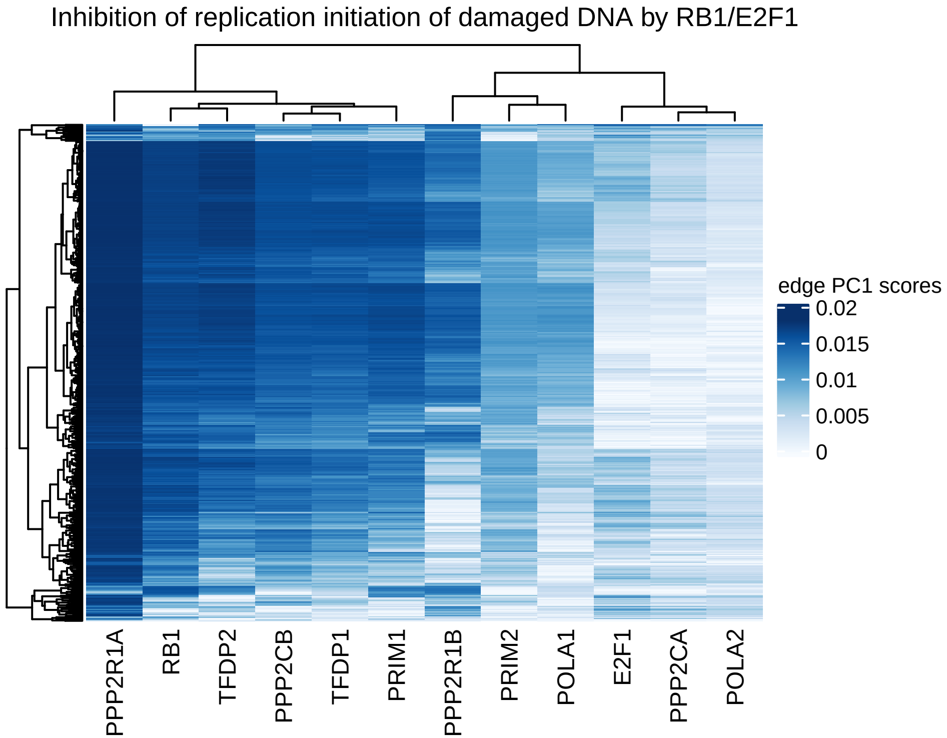

Inhibition of replication initiation of damaged DNA by RB1/E2F1
edge PC1 scores
0.02
0.015
0.01
0.005
0
PPP2R1A
RB1
TFDP2
PPP2CB
TFDP1
PRIM1
PPP2R1B
PRIM2
POLA1
E2F1
PPP2CA
POLA2

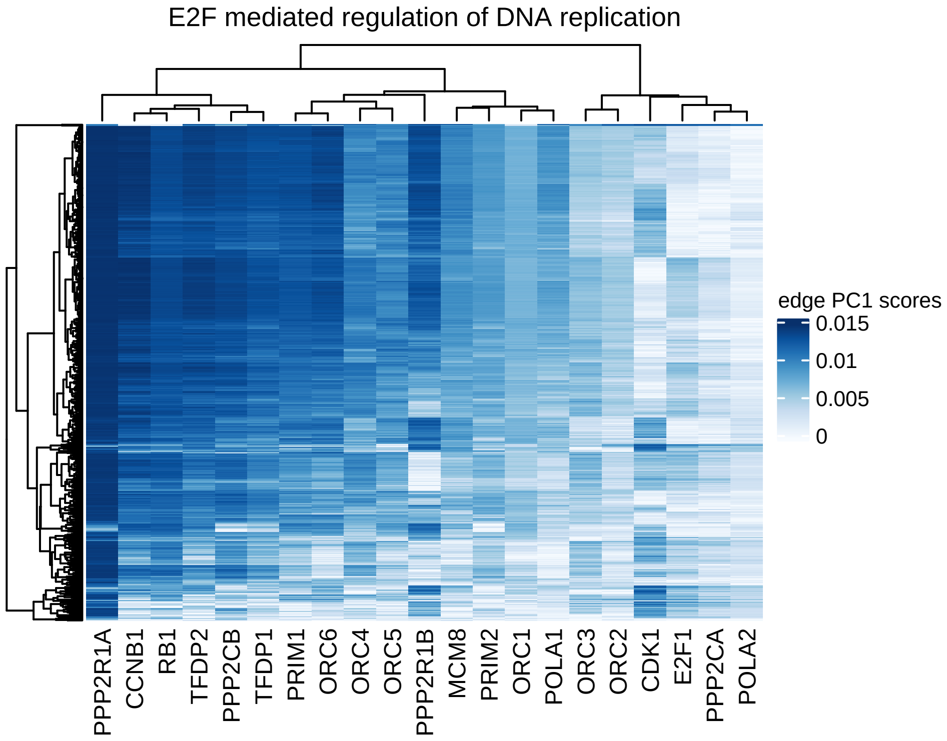

E2F mediated regulation of DNA replication
edge PC1 scores
0.015
0.01
0.005
0
PPP2R1A
CCNB1
RB1
TFDP2
PPP2CB
TFDP1
PRIM1
ORC6
ORC4
ORC5
PPP2R1B
MCM8
PRIM2
ORC1
POLA1
ORC3
ORC2
CDK1
E2F1
PPP2CA
POLA2

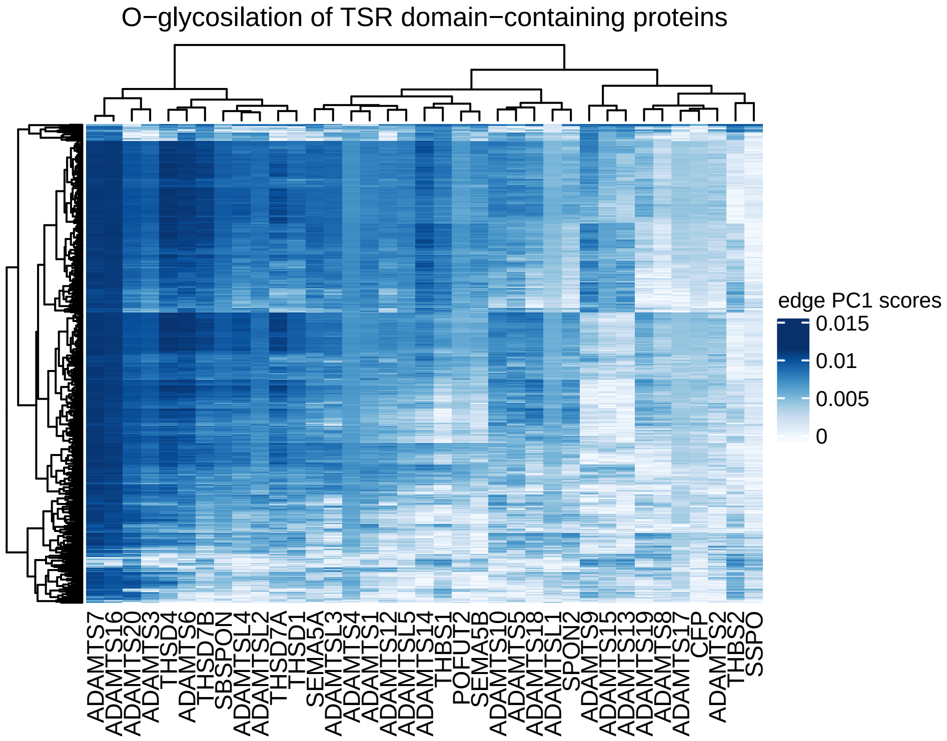
O−glycosilation of TSR domain−containing proteins
edge PC1 scores
0.015
0.01
0.005
0
ADAMTS7
ADAMTS16
ADAMTS20
ADAMTS3
THSD4
ADAMTS6
THSD7B
SBSPON
ADAMTSL4
ADAMTSL2
THSD7A
THSD1
SEMA5A
ADAMTSL3
ADAMTS4
ADAMTS1
ADAMTS12
ADAMTSL5
ADAMTS14
THBS1
POFUT2
SEMA5B
ADAMTS10
ADAMTS5
ADAMTS18
ADAMTSL1
SPON2
ADAMTS9
ADAMTS15
ADAMTS13
ADAMTS19
ADAMTS8
ADAMTS17
CFP
ADAMTS2
THBS2
SSPO

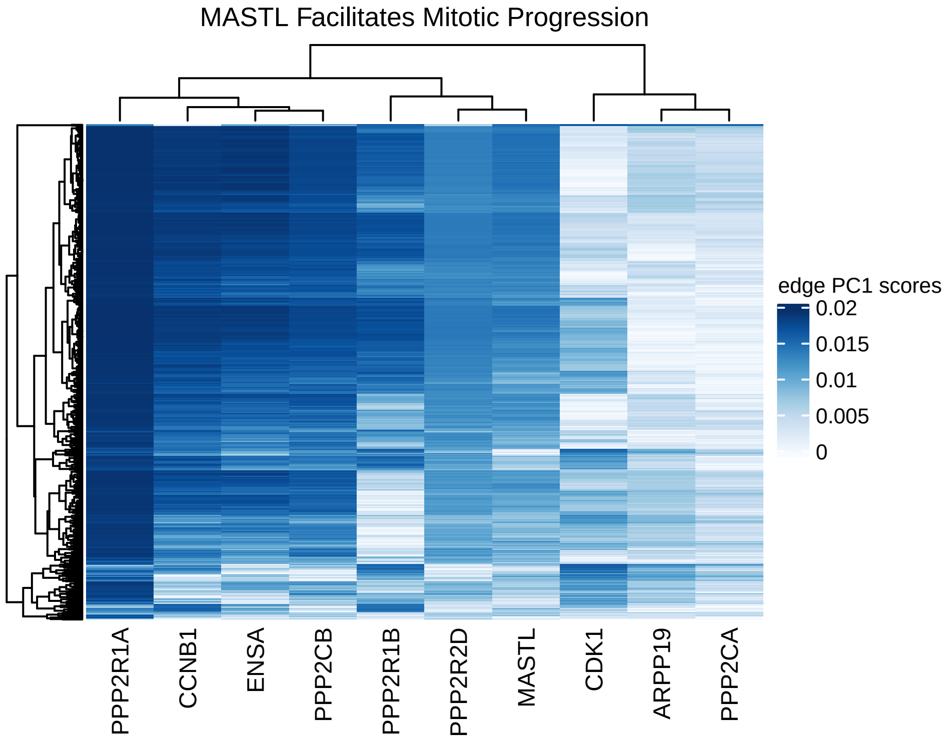

MASTL Facilitates Mitotic Progression
edge PC1 scores
0.02
0.015
0.01
0.005
0
PPP2R1A
CCNB1
ENSA
PPP2CB
PPP2R1B
PPP2R2D
MASTL
CDK1
ARPP19
PPP2CA

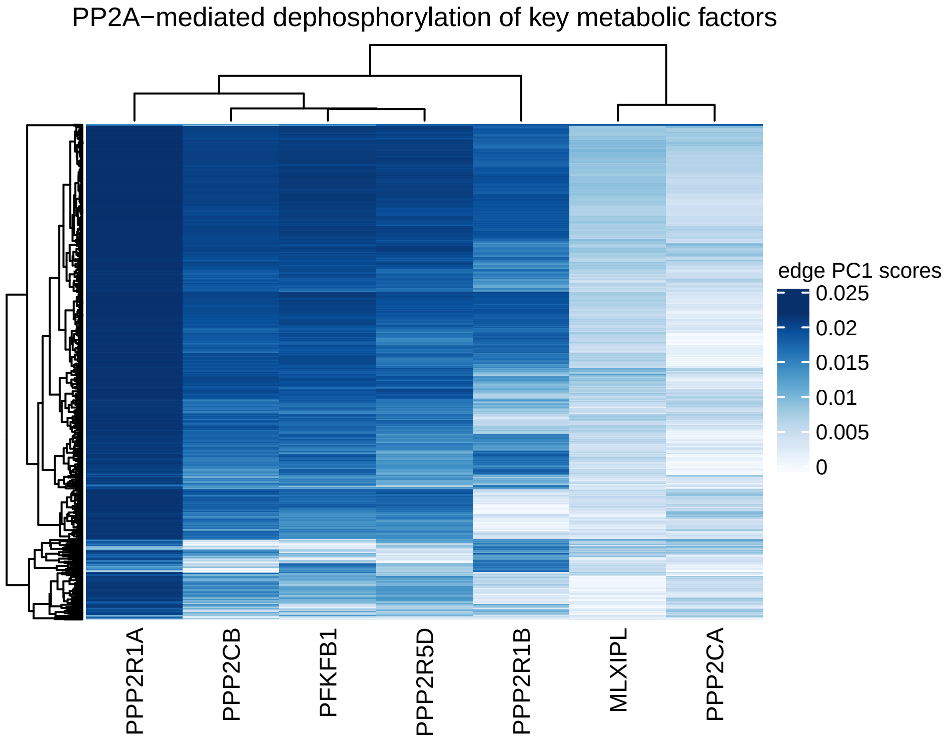
PP2A−mediated dephosphorylation of key metabolic factors
edge PC1 scores
0.025
0.02
0.015
0.01
0.005
0
PPP2R1A
PPP2CB
PFKFB1
PPP2R5D
PPP2R1B
MLXIPL
PPP2CA

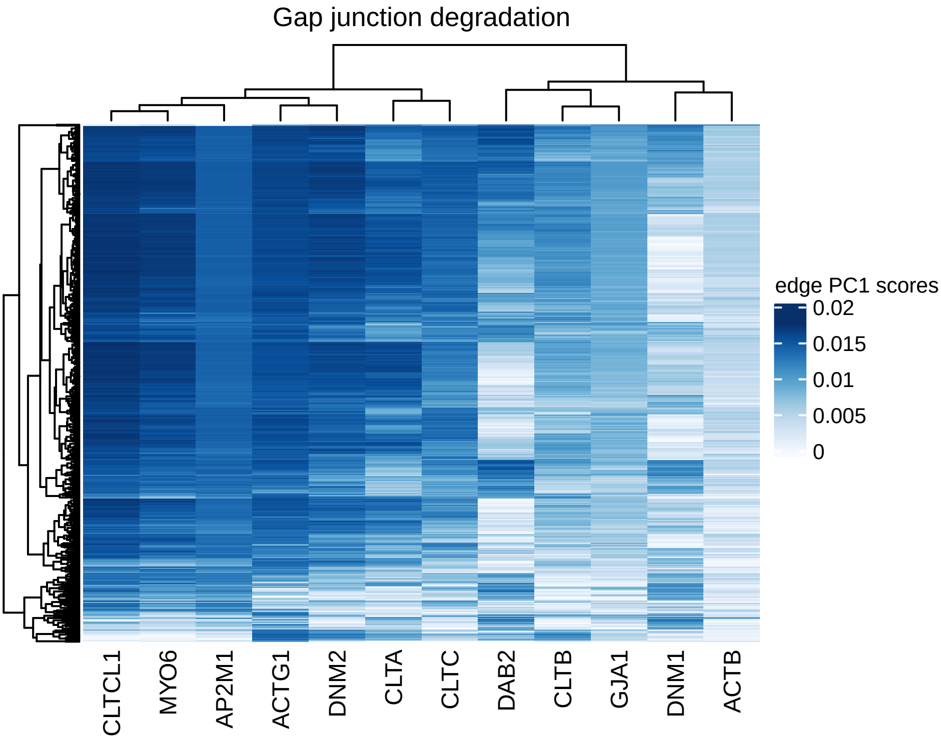
Gap junction degradation
edge PC1 scores
0.02
0.015
0.01
0.005
0
CLTCL1
MYO6
AP2M1
ACTG1
DNM2
CLTA
CLTC
DAB2
CLTB
GJA1
DNM1
ACTB

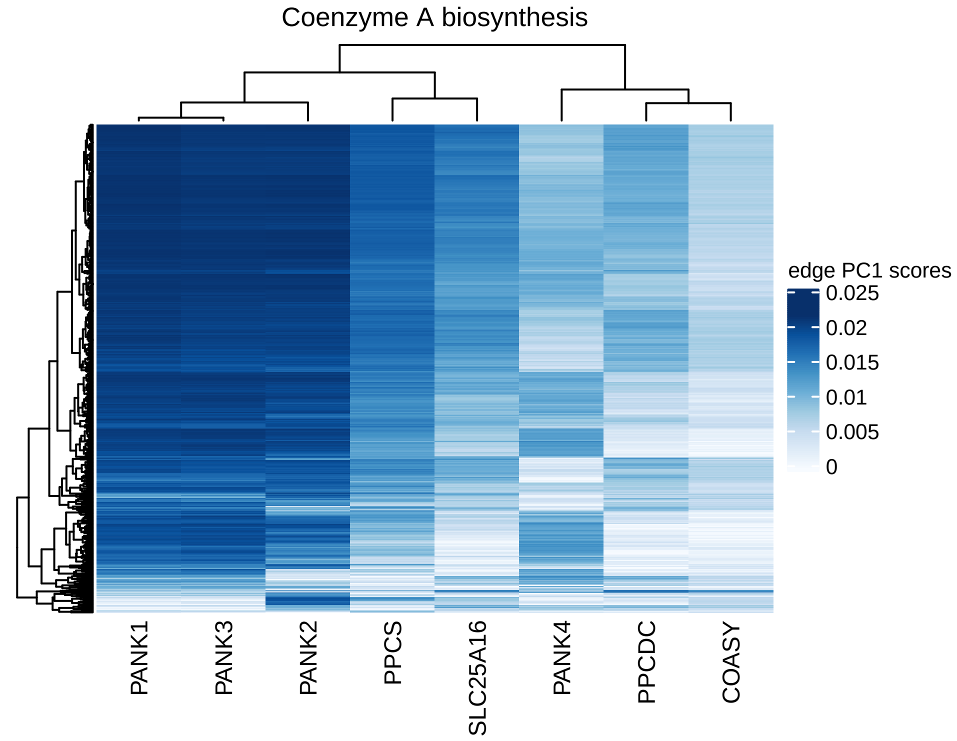

Coenzyme A biosynthesis
edge PC1 scores
0.025
0.02
0.015
0.01
0.005
0
PANK1
PANK3
PANK2
PPCS
SLC25A16
PANK4
PPCDC
COASY

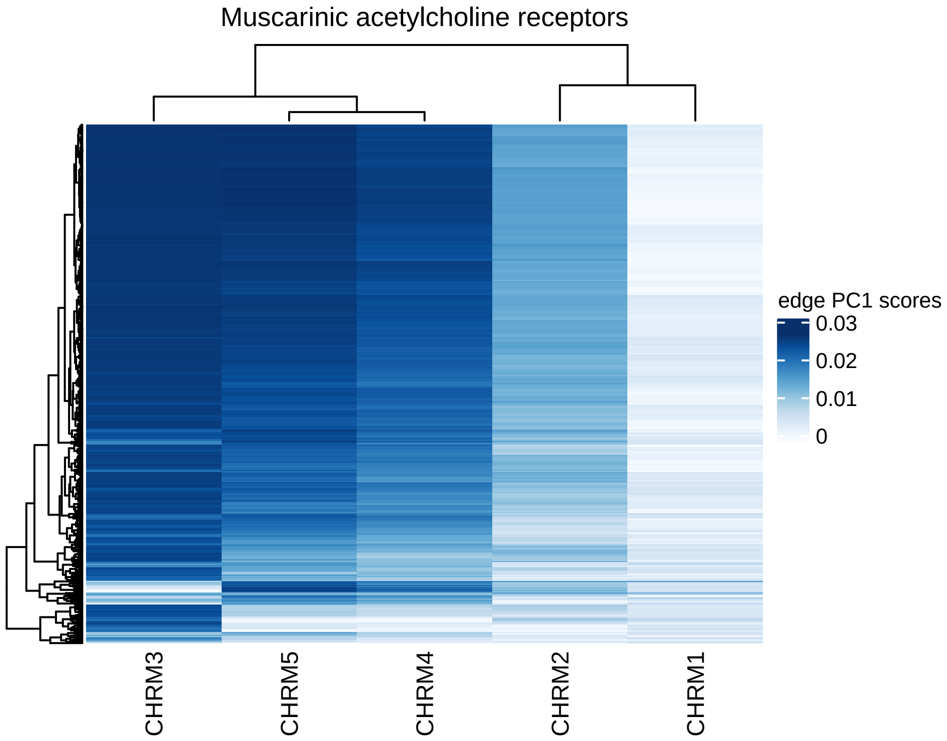

Muscarinic acetylcholine receptors
edge PC1 scores
0.03
0.02
0.01
0
CHRM3
CHRM5
CHRM4
CHRM2
CHRM1

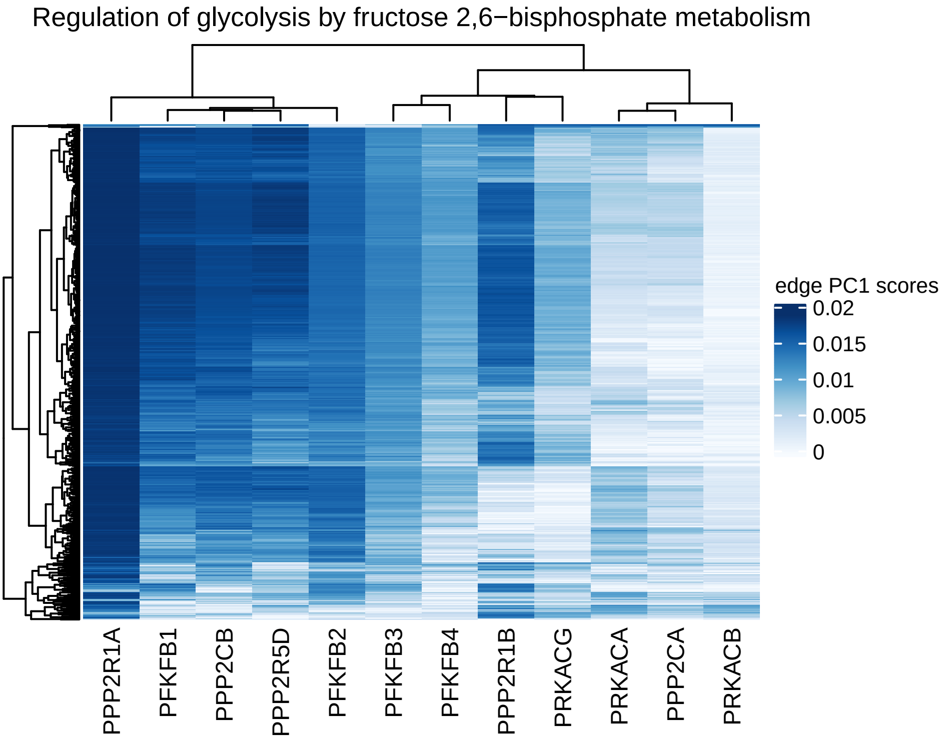

Regulation of glycolysis by fructose 2,6−bisphosphate metabolism
edge PC1 scores
0.02
0.015
0.01
0.005
0
PPP2R1A
PFKFB1
PPP2CB
PPP2R5D
PFKFB2
PFKFB3
PFKFB4
PPP2R1B
PRKACG
PRKACA
PPP2CA
PRKACB

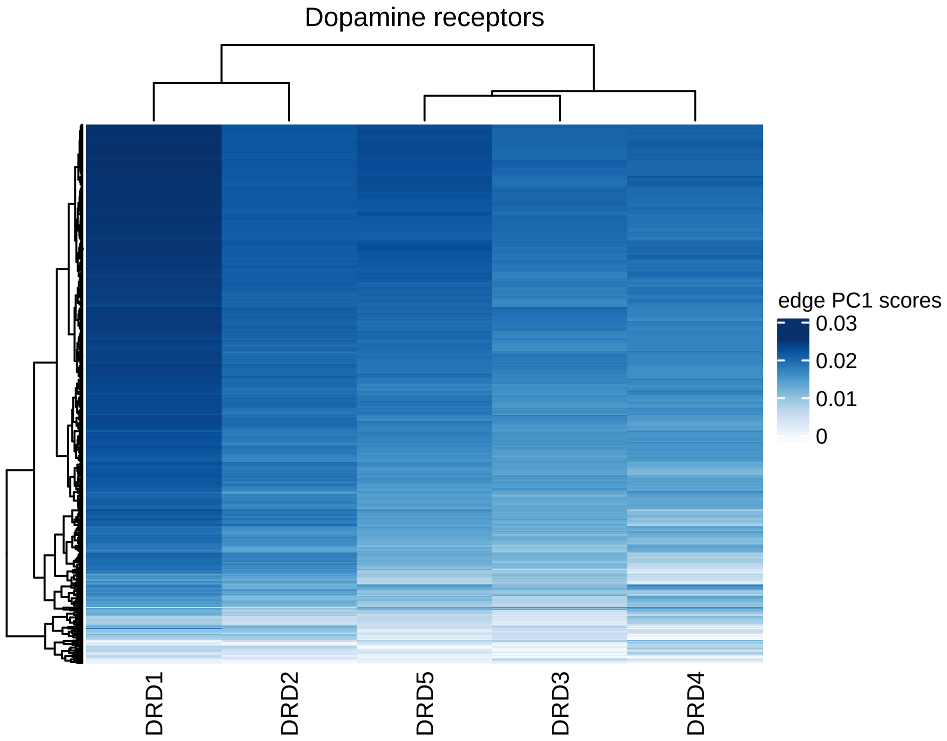
Dopamine receptors
edge PC1 scores
0.03
0.02
0.01
0
DRD1
DRD2
DRD5
DRD3
DRD4

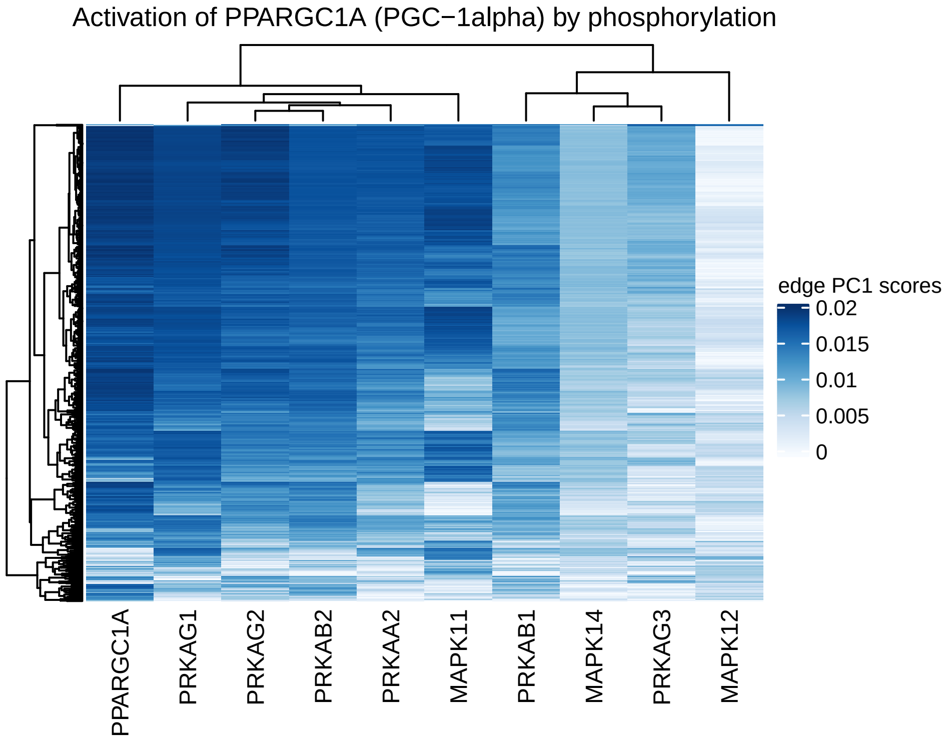
Activation of PPARGC1A (PGC−1alpha) by phosphorylation
edge PC1 scores
0.02
0.015
0.01
0.005
0
PPARGC1A
PRKAG1
PRKAG2
PRKAB2
PRKAA2
MAPK11
PRKAB1
MAPK14
PRKAG3
MAPK12

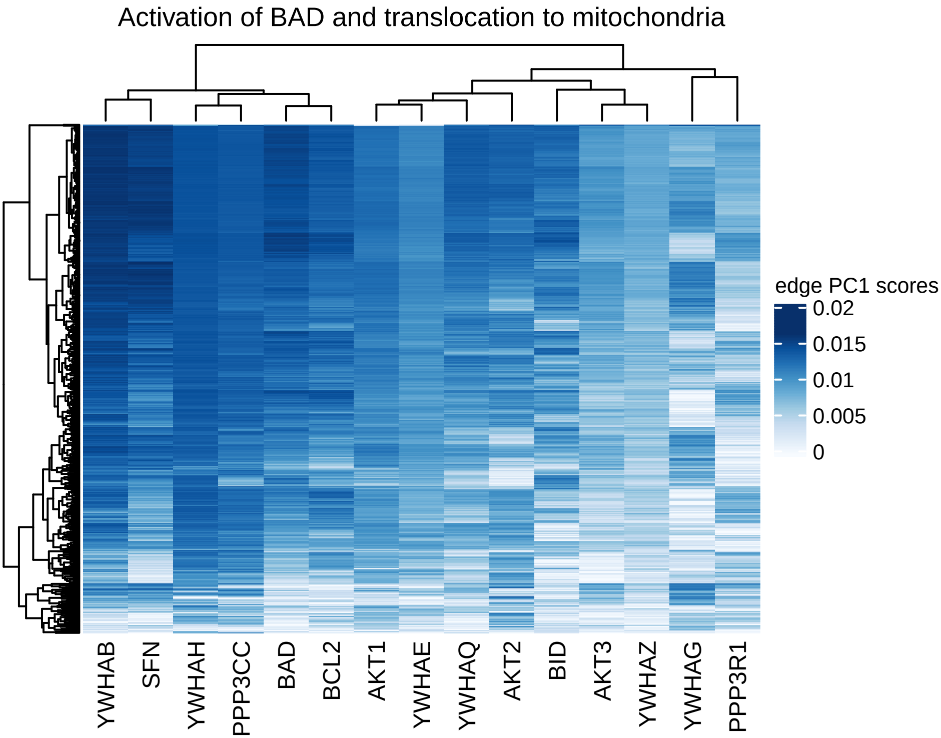
Activation of BAD and translocation to mitochondria
edge PC1 scores
0.02
0.015
0.01
0.005
0
YWHAB
SFN
YWHAH
PPP3CC
BAD
BCL2
AKT1
YWHAE
YWHAQ
AKT2
BID
AKT3
YWHAZ
YWHAG
PPP3R1

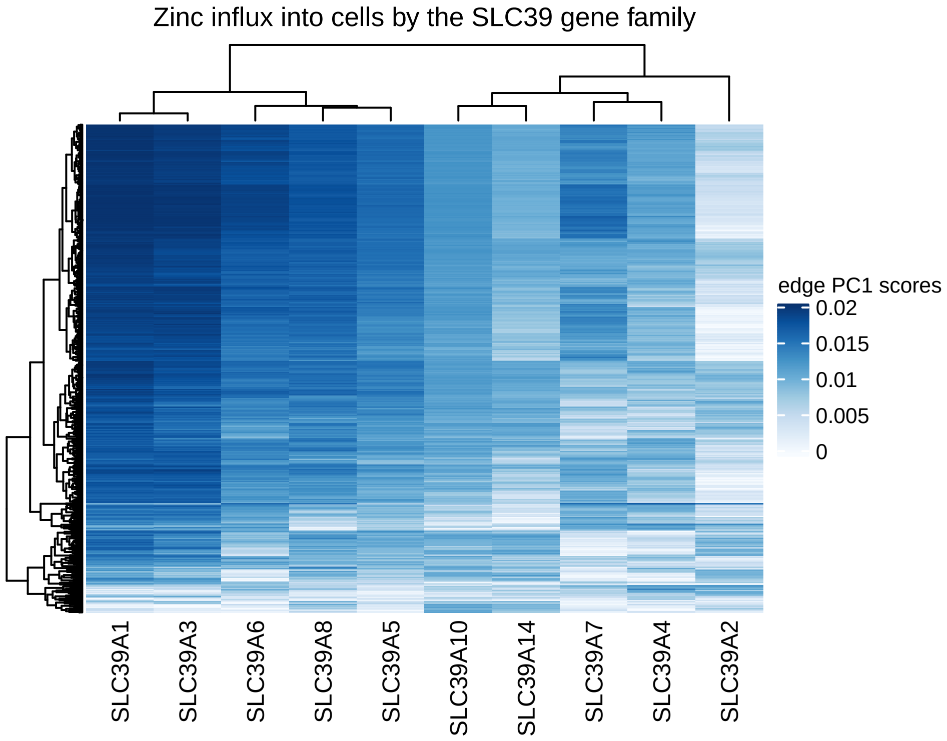
Zinc influx into cells by the SLC39 gene family
edge PC1 scores
0.02
0.015
0.01
0.005
0
SLC39A1
SLC39A3
SLC39A6
SLC39A8
SLC39A5
SLC39A10
SLC39A14
SLC39A7
SLC39A4
SLC39A2

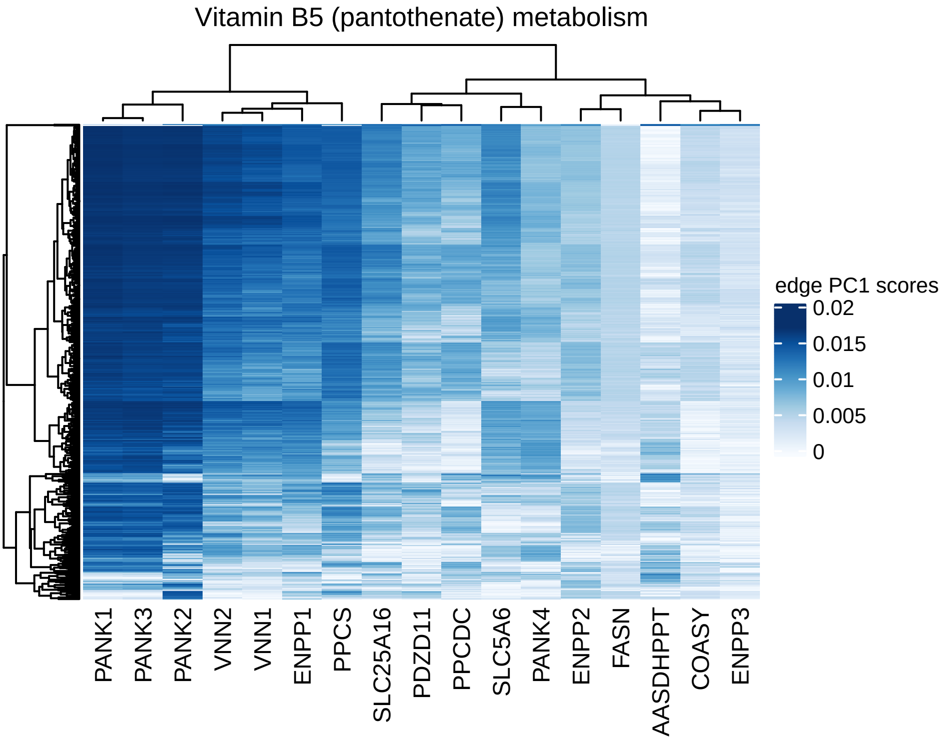
Vitamin B5 (pantothenate) metabolism
edge PC1 scores
0.02
0.015
0.01
0.005
0
PANK1
PANK3
PANK2
VNN2
VNN1
ENPP1
PPCS
SLC25A16
PDZD11
PPCDC
SLC5A6
PANK4
ENPP2
FASN
AASDHPPT
COASY
ENPP3

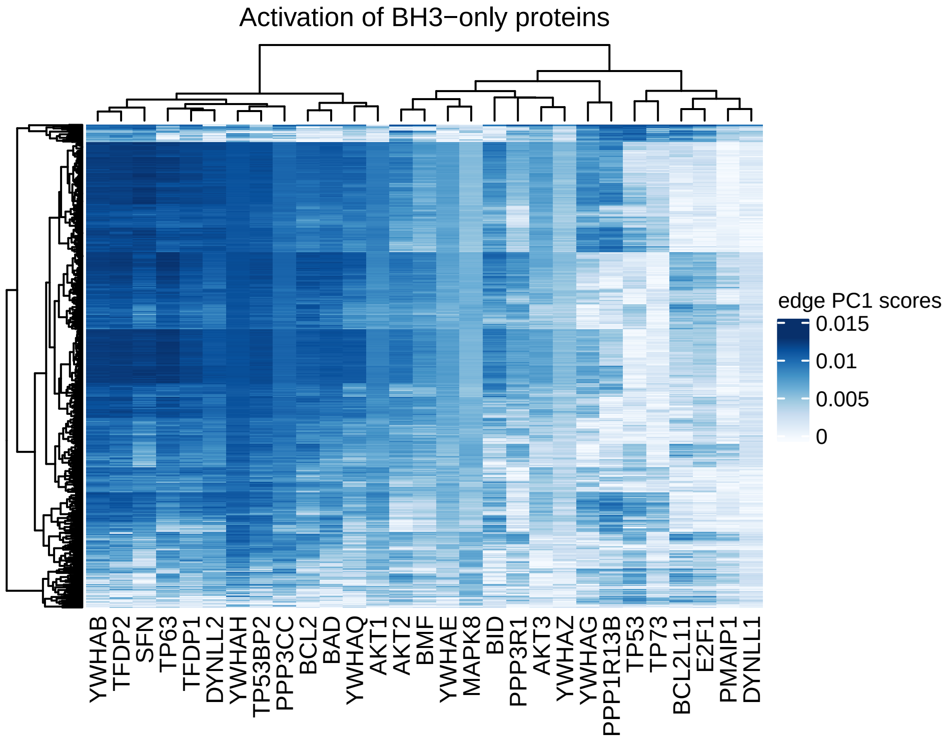
Activation of BH3−only proteins
edge PC1 scores
0.015
0.01
0.005
0
YWHAB
TFDP2
SFN
TP63
TFDP1
DYNLL2
YWHAH
TP53BP2
PPP3CC
BCL2
BAD
YWHAQ
AKT1
AKT2
BMF
YWHAE
MAPK8
BID
PPP3R1
AKT3
YWHAZ
YWHAG
PPP1R13B
TP53
TP73
BCL2L11
E2F1
PMAIP1
DYNLL1

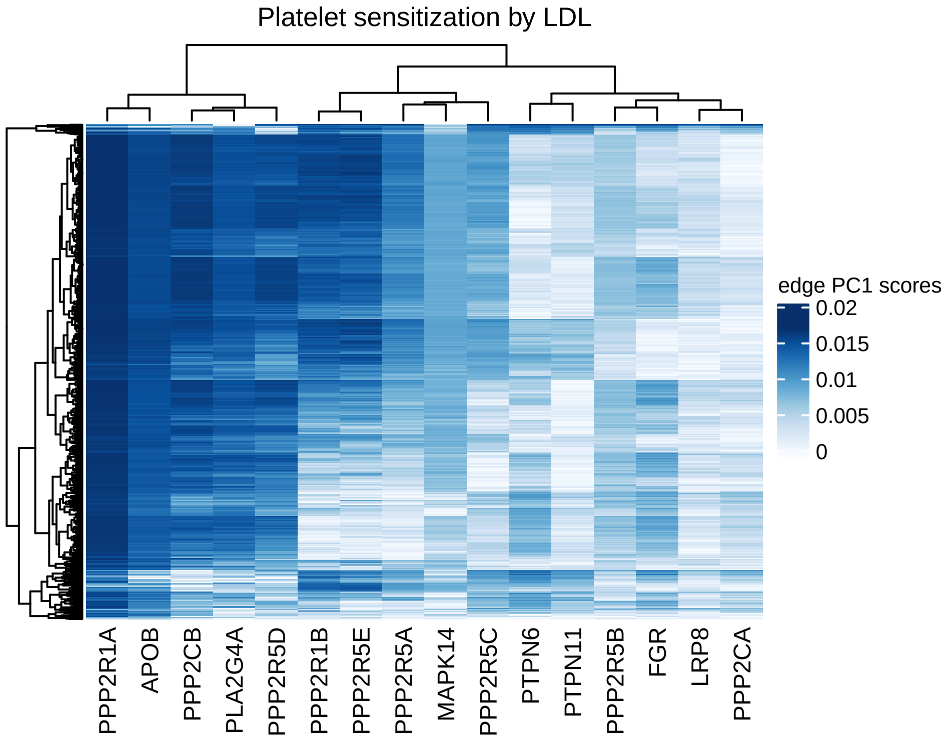

Platelet sensitization by LDL
edge PC1 scores
0.02
0.015
0.01
0.005
0
PPP2R1A
APOB
PPP2CB
PLA2G4A
PPP2R5D
PPP2R1B
PPP2R5E
PPP2R5A
MAPK14
PPP2R5C
PTPN6
PTPN11
PPP2R5B
FGR
LRP8
PPP2CA

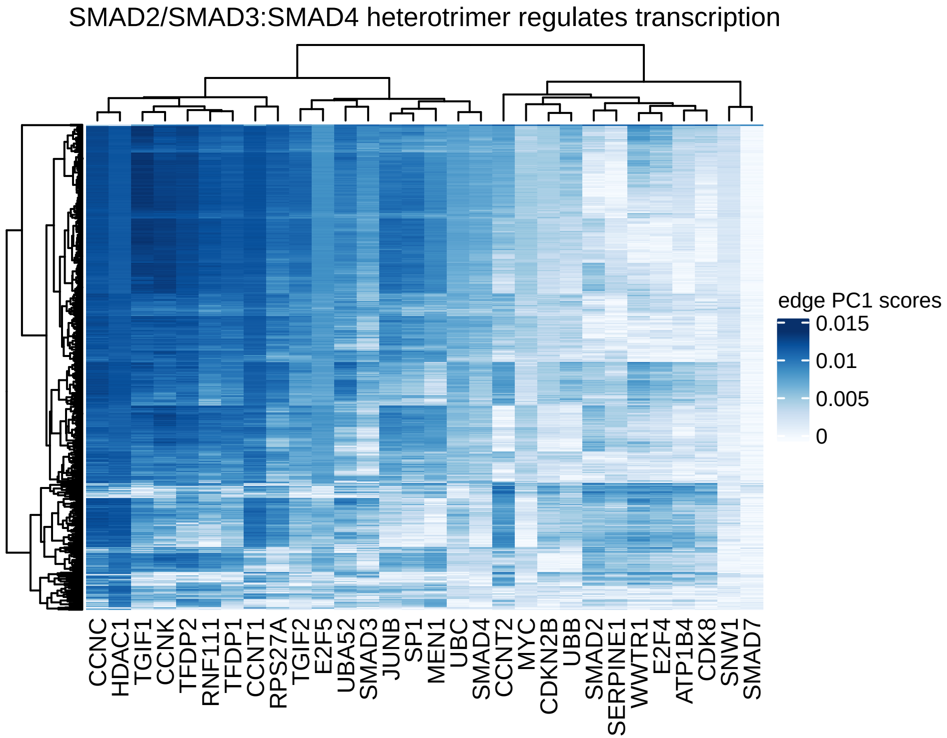

SMAD2/SMAD3:SMAD4 heterotrimer regulates transcription
edge PC1 scores
0.015
0.01
0.005
0
CCNC
HDAC1
TGIF1
CCNK
TFDP2
RNF111
TFDP1
CCNT1
RPS27A
TGIF2
E2F5
UBA52
SMAD3
JUNB
SP1
MEN1
UBC
SMAD4
CCNT2
MYC
CDKN2B
UBB
SMAD2
SERPINE1
WWTR1
E2F4
ATP1B4
CDK8
SNW1
SMAD7

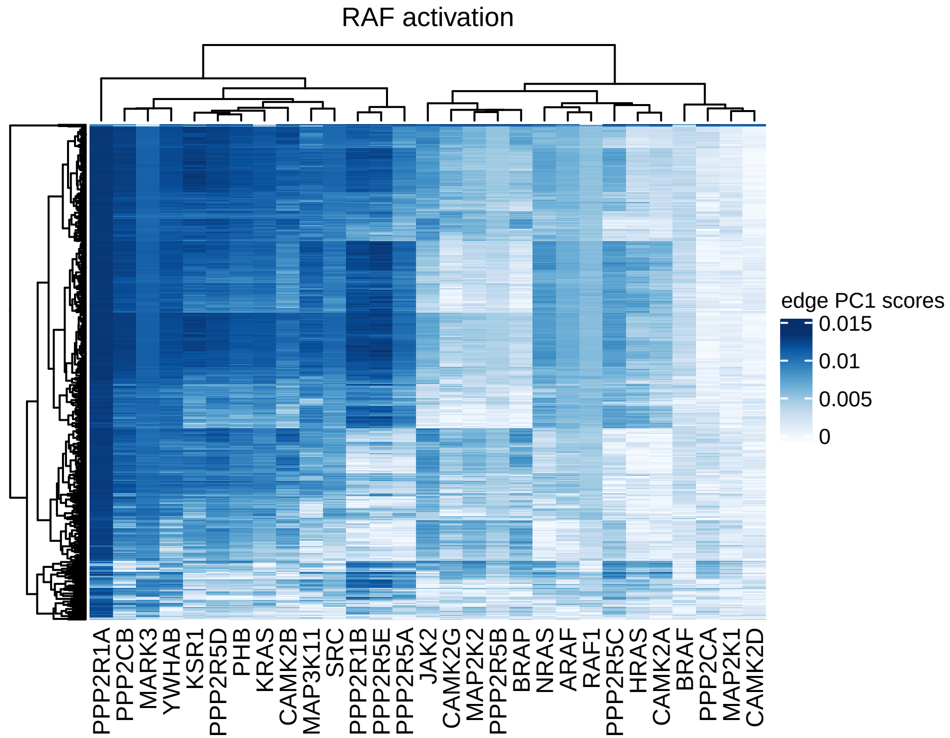

RAF activation
edge PC1 scores
0.015
0.01
0.005
0
PPP2R1A
PPP2CB
MARK3
YWHAB
KSR1
PPP2R5D
PHB
KRAS
CAMK2B
MAP3K11
SRC
PPP2R1B
PPP2R5E
PPP2R5A
JAK2
CAMK2G
MAP2K2
PPP2R5B
BRAP
NRAS
ARAF
RAF1
PPP2R5C
HRAS
CAMK2A
BRAF
PPP2CA
MAP2K1
CAMK2D

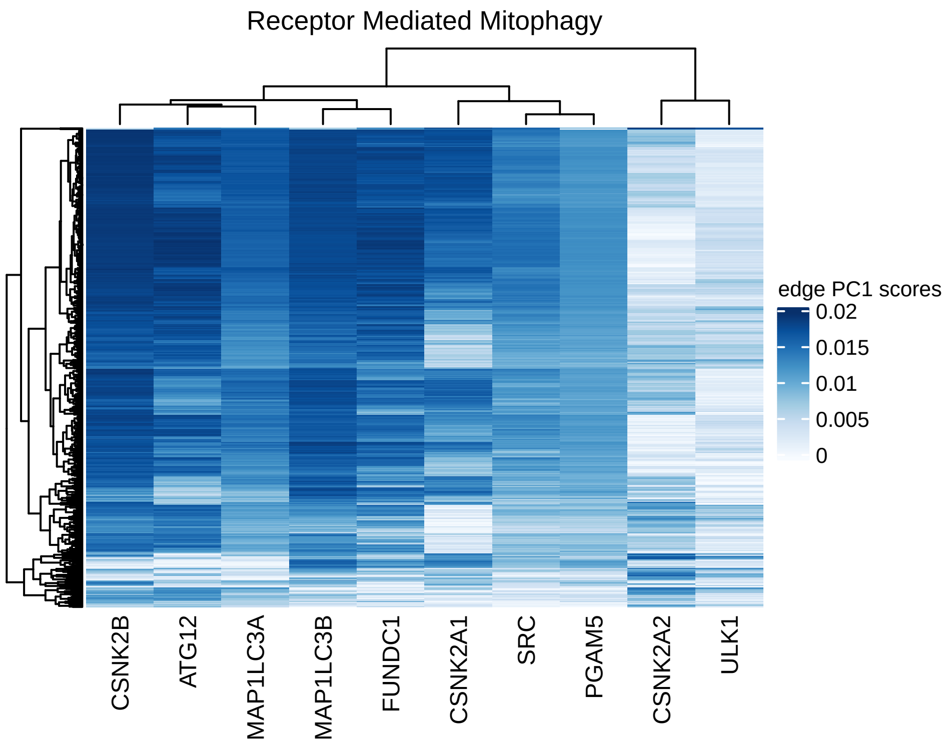
Receptor Mediated Mitophagy
edge PC1 scores
0.02
0.015
0.01
0.005
0
CSNK2B
ATG12
MAP1LC3A
MAP1LC3B
FUNDC1
CSNK2A1
SRC
PGAM5
CSNK2A2
ULK1

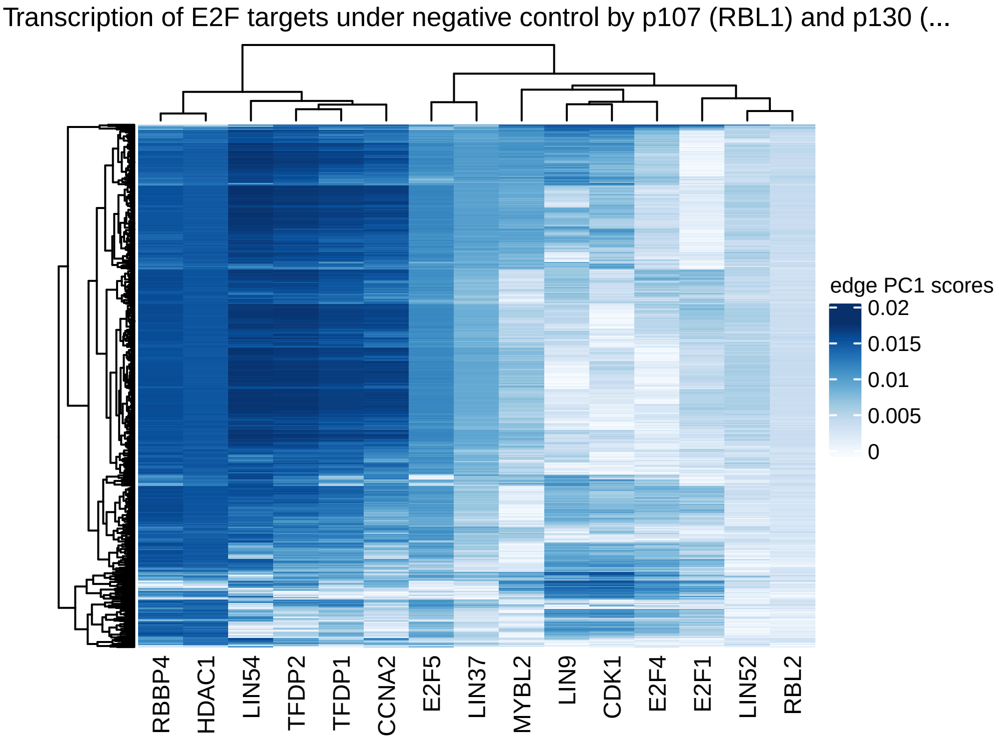

Transcription of E2F targets under negative control by p107 (RBL1) and p130 (...
edge PC1 scores
0.02
0.015
0.01
0.005
0
RBBP4
HDAC1
LIN54
TFDP2
TFDP1
CCNA2
E2F5
LIN37
MYBL2
LIN9
CDK1
E2F4
E2F1
LIN52
RBL2

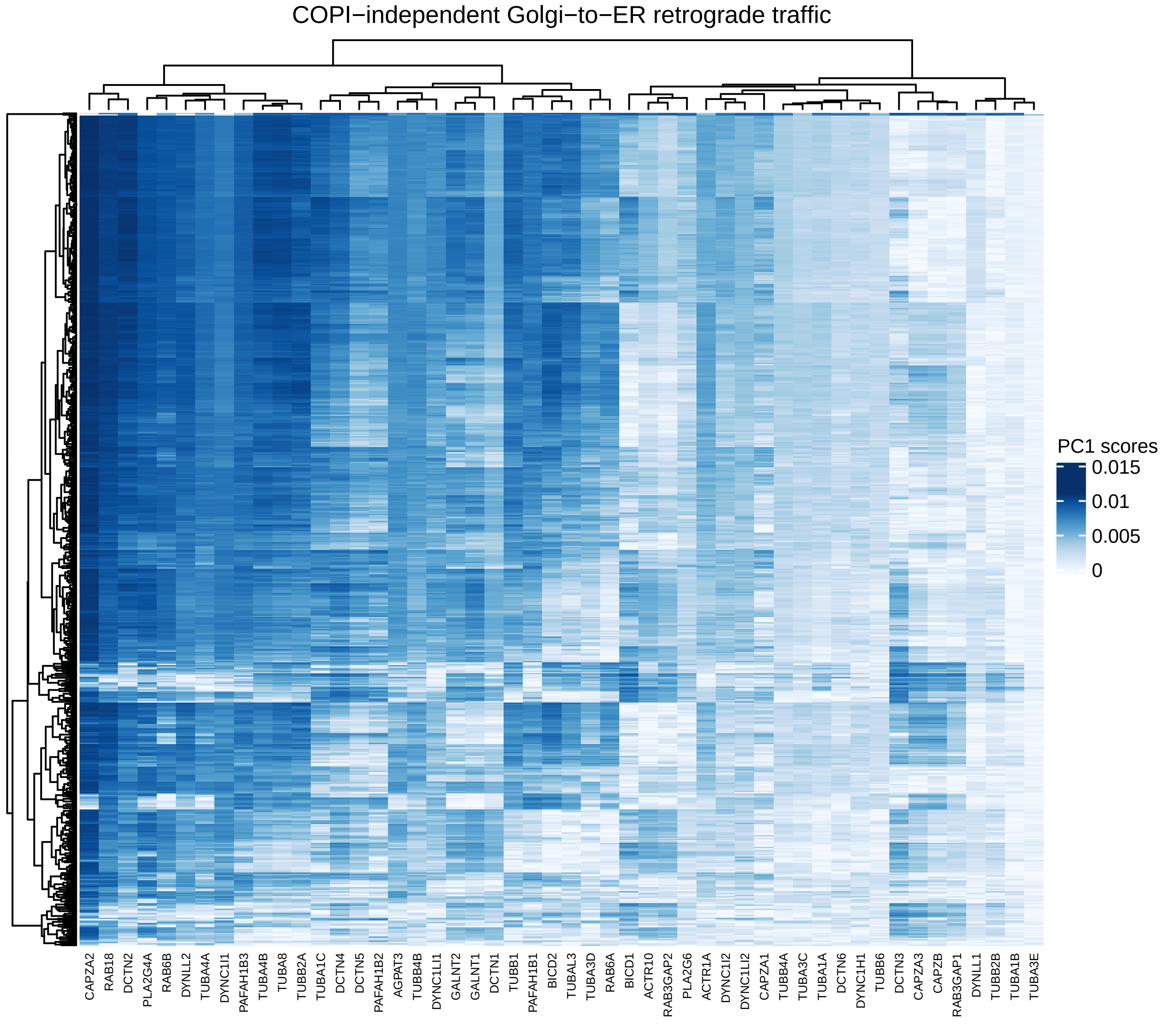

COPI−independent Golgi−to−ER retrograde traffic
PC1 scores
0.015
0.01
0.005
0
CAPZA2
RAB18
DCTN2
PLA2G4A
RAB6B
DYNLL2
TUBA4A
DYNC1I1
PAFAH1B3
TUBA4B
TUBA8
TUBB2A
TUBA1C
DCTN4
DCTN5
PAFAH1B2
AGPAT3
TUBB4B
DYNC1LI1
GALNT2
GALNT1
DCTN1
TUBB1
PAFAH1B1
BICD2
TUBAL3
TUBA3D
RAB6A
BICD1
ACTR10
RAB3GAP2
PLA2G6
ACTR1A
DYNC1I2
DYNC1LI2
CAPZA1
TUBB4A
TUBA3C
TUBA1A
DCTN6
DYNC1H1
TUBB6
DCTN3
CAPZA3
CAPZB
RAB3GAP1
DYNLL1
TUBB2B
TUBA1B
TUBA3E

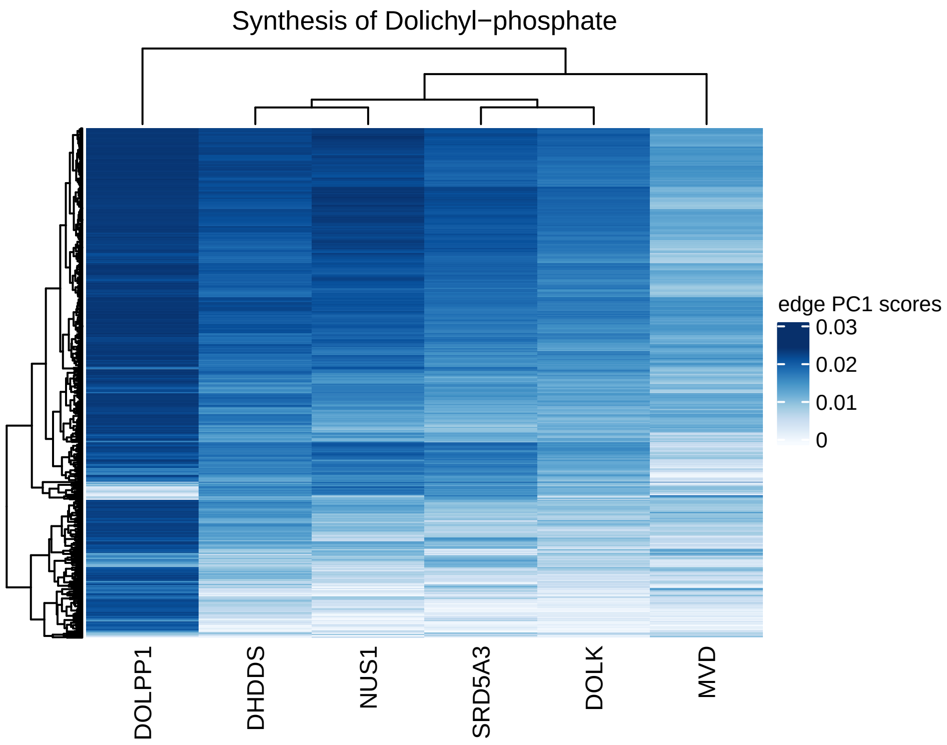

Synthesis of Dolichyl−phosphate
edge PC1 scores
0.03
0.02
0.01
0
DOLPP1
DHDDS
NUS1
SRD5A3
DOLK
MVD

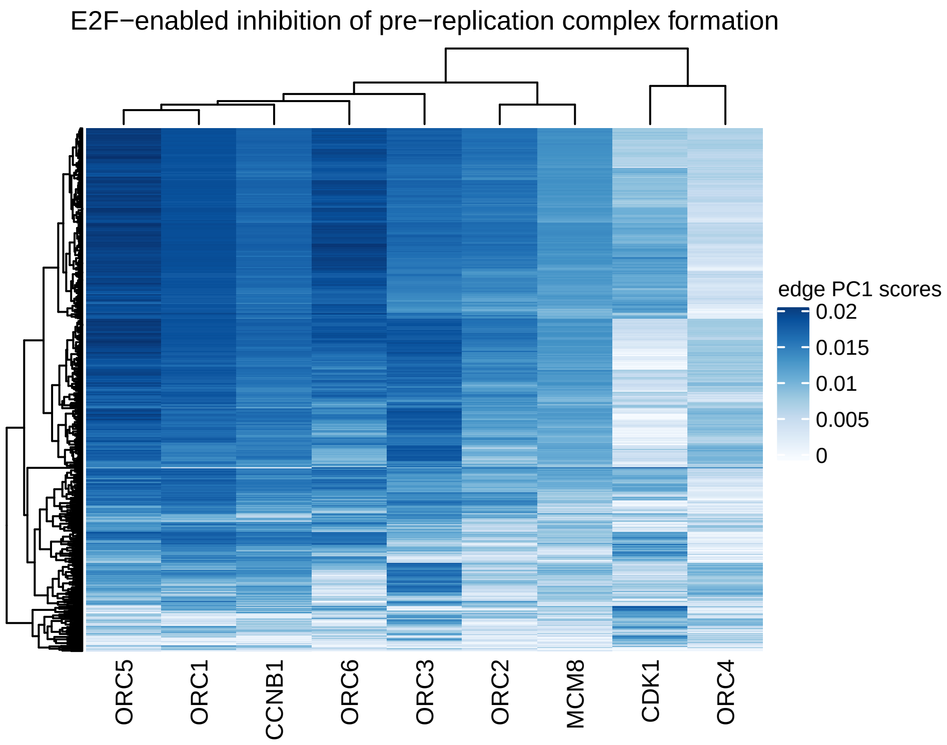
E2F−enabled inhibition of pre−replication complex formation
edge PC1 scores
0.02
0.015
0.01
0.005
0
ORC5
ORC1
CCNB1
ORC6
ORC3
ORC2
MCM8
CDK1
ORC4

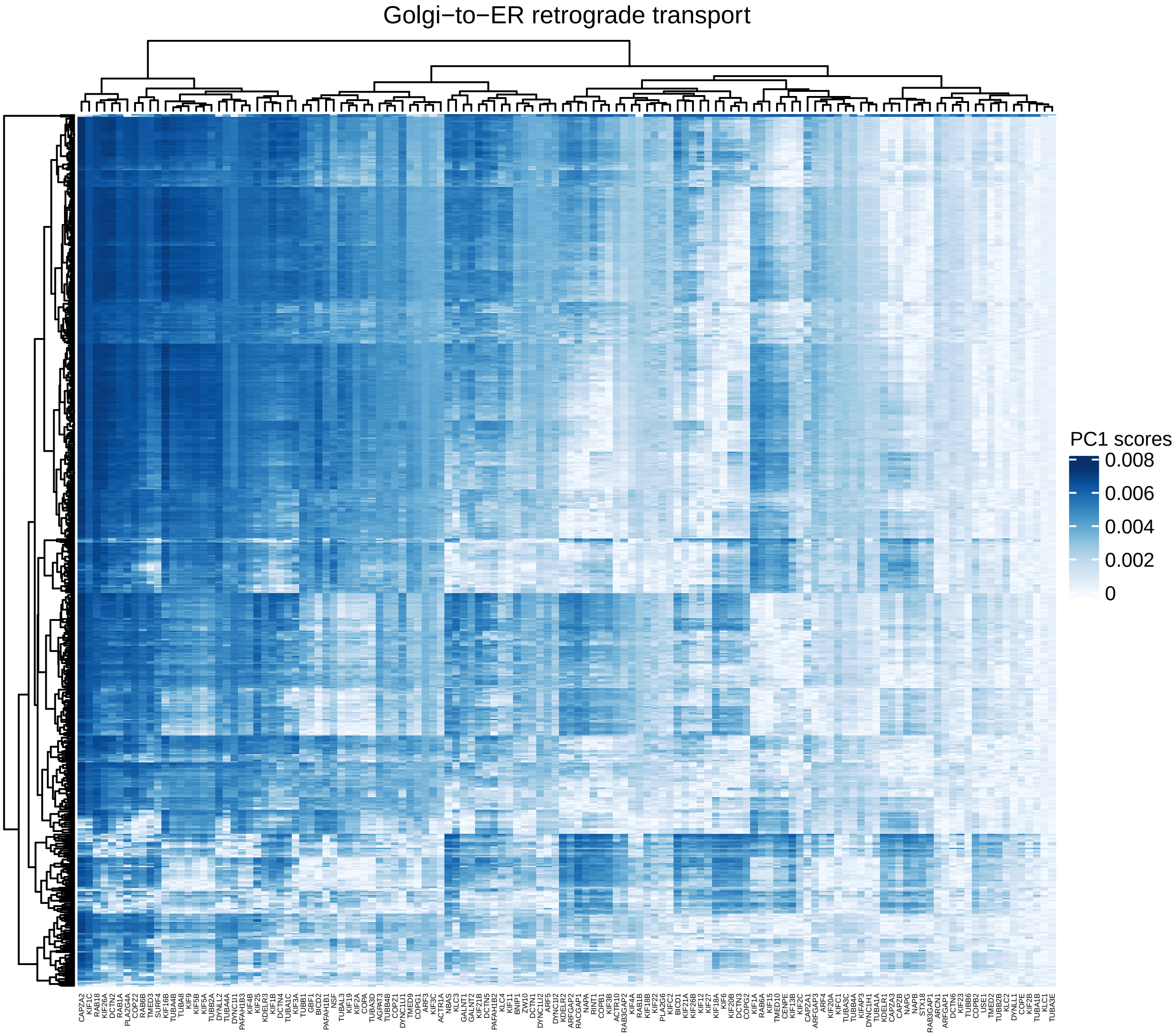
Golgi−to−ER retrograde transport
PC1 scores
0.008
0.006
0.004
0.002
0

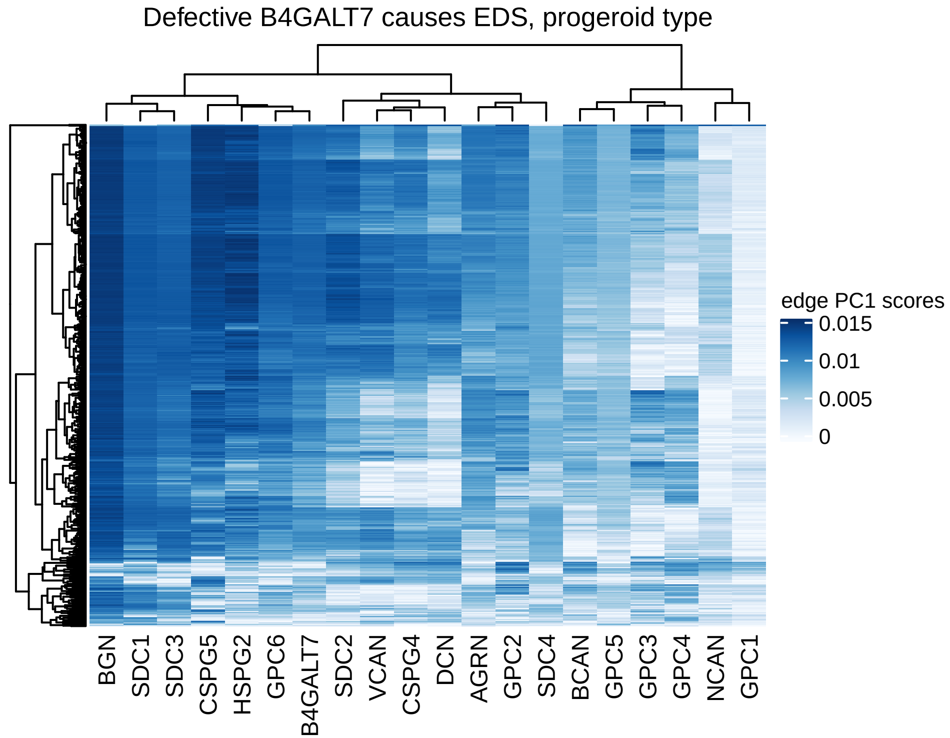
Defective B4GALT7 causes EDS, progeroid type
BGN
SDC1
SDC3
CSPG5
HSPG2
GPC6
B4GALT7
SDC2
VCAN
CSPG4
DCN
AGRN
GPC2
SDC4
BCAN
GPC5
GPC3
GPC4
NCAN
GPC1
edge PC1 scores
0.015
0.01
0.005
0

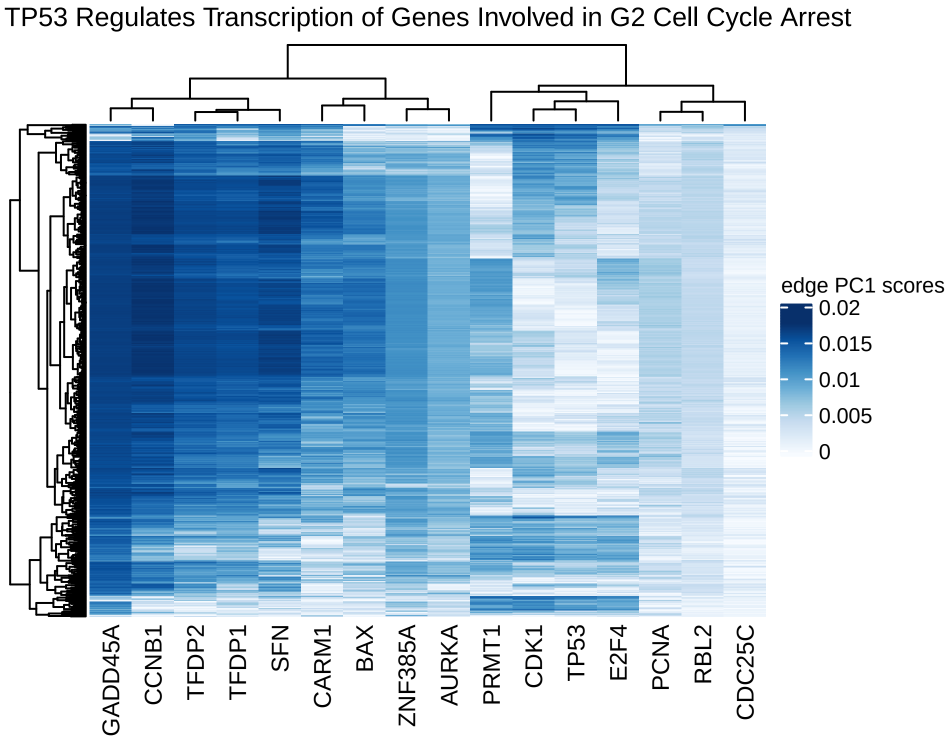
TP53 Regulates Transcription of Genes Involved in G2 Cell Cycle Arrest
edge PC1 scores
0.02
0.015
0.01
0.005
0
GADD45A
CCNB1
TFDP2
TFDP1
SFN
CARM1
BAX
ZNF385A
AURKA
PRMT1
CDK1
TP53
E2F4
PCNA
RBL2
CDC25C

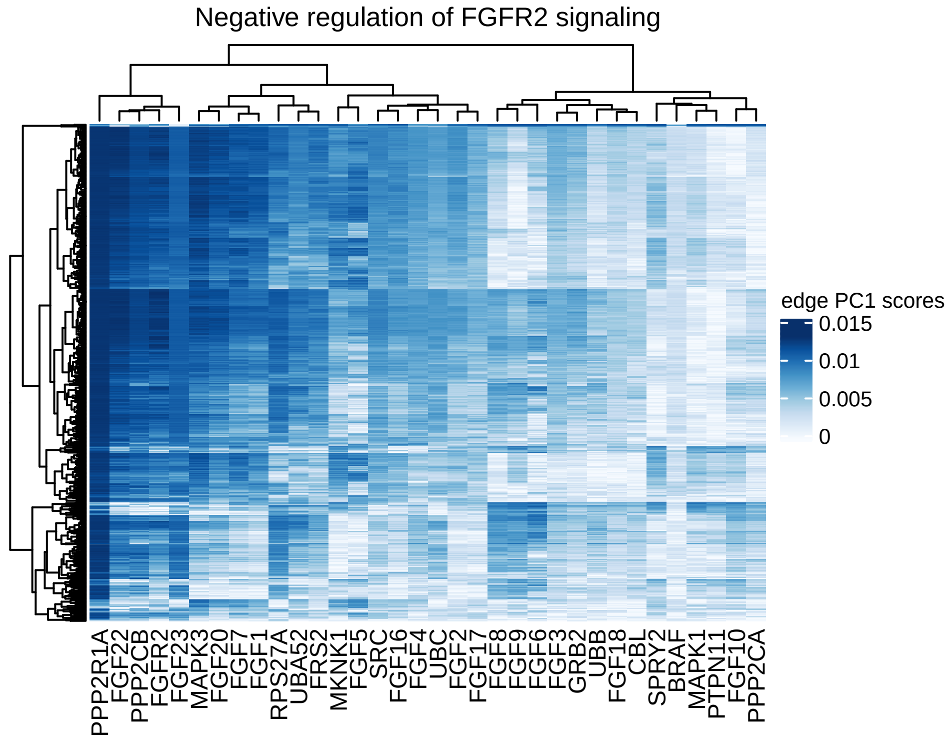
Negative regulation of FGFR2 signaling
edge PC1 scores
0.015
0.01
0.005
0
PPP2R1A
FGF22
PPP2CB
FGFR2
FGF23
MAPK3
FGF20
FGF7
FGF1
RPS27A
UBA52
FRS2
MKNK1
FGF5
SRC
FGF16
FGF4
UBC
FGF2
FGF17
FGF8
FGF9
FGF6
FGF3
GRB2
UBB
FGF18
CBL
SPRY2
BRAF
MAPK1
PTPN11
FGF10
PPP2CA

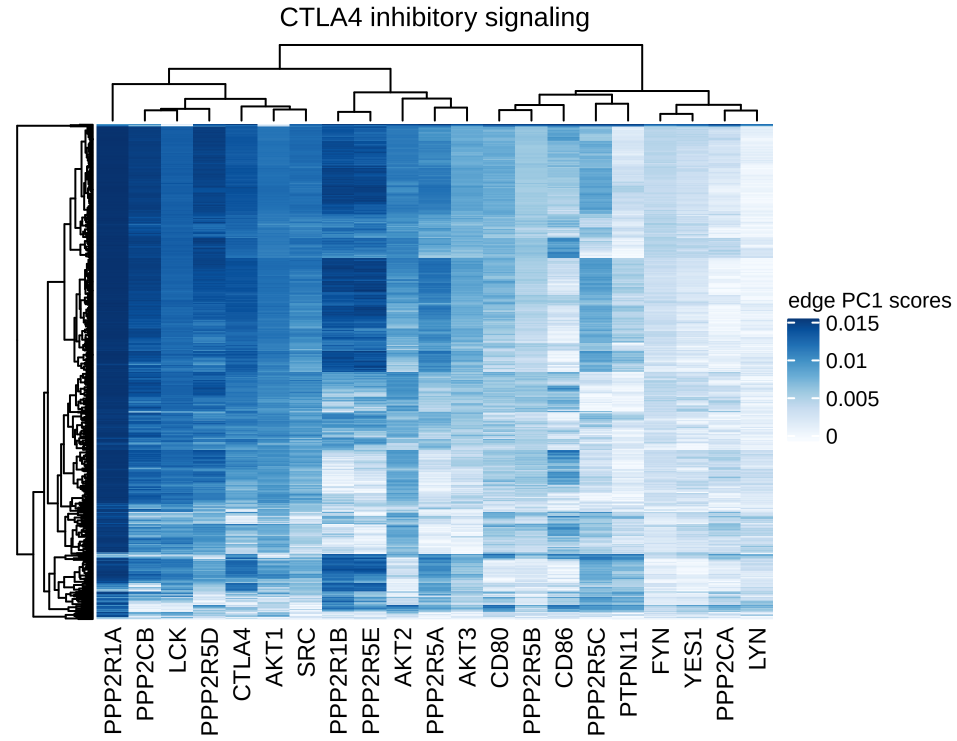
CTLA4 inhibitory signaling
edge PC1 scores
0.015
0.01
0.005
0
PPP2R1A
PPP2CB
LCK
PPP2R5D
CTLA4
AKT1
SRC
PPP2R1B
PPP2R5E
AKT2
PPP2R5A
AKT3
CD80
PPP2R5B
CD86
PPP2R5C
PTPN11
FYN
YES1
PPP2CA
LYN

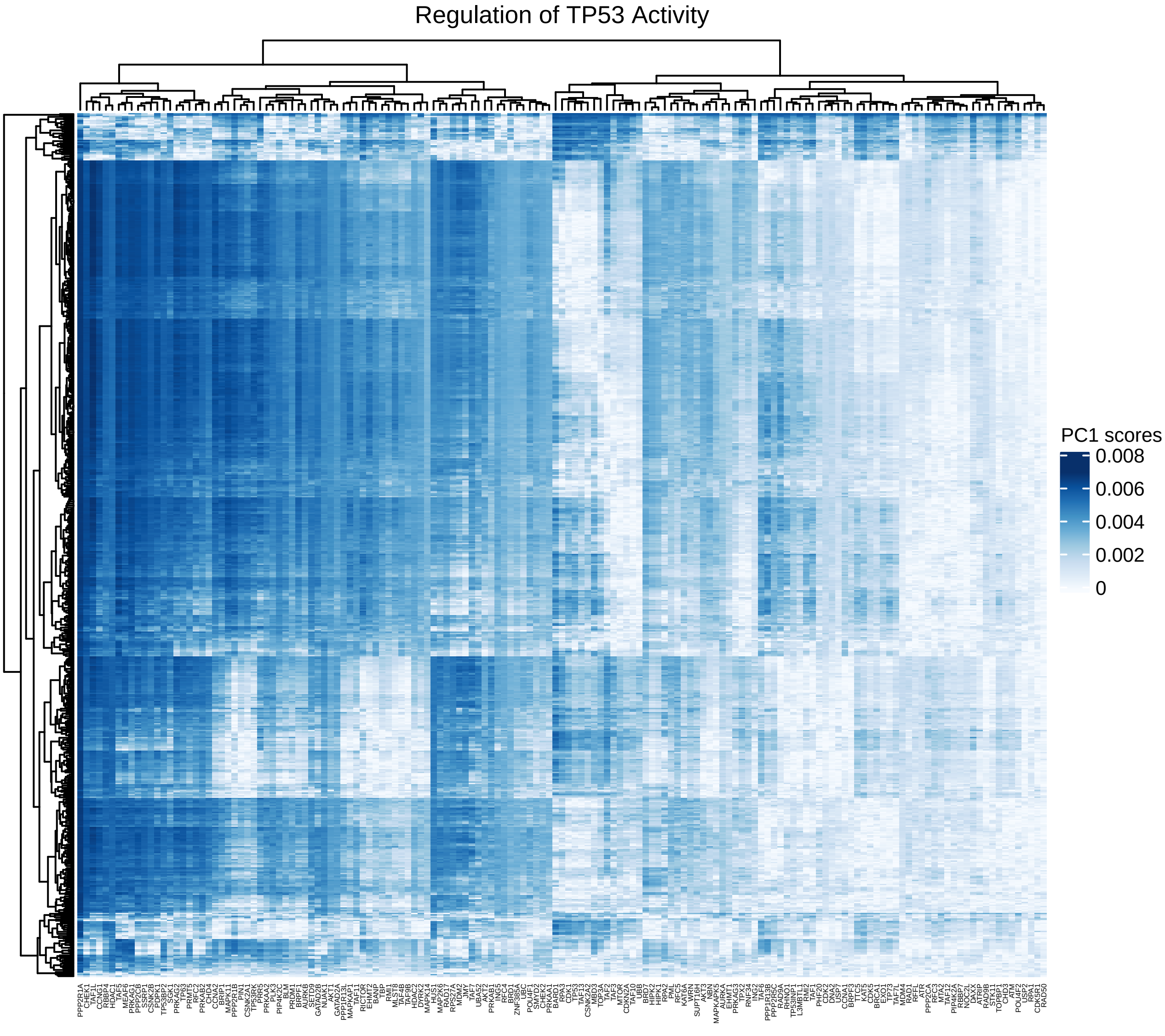
Regulation of TP53 Activity
PC1 scores
0.008
0.006
0.004
0.002
0

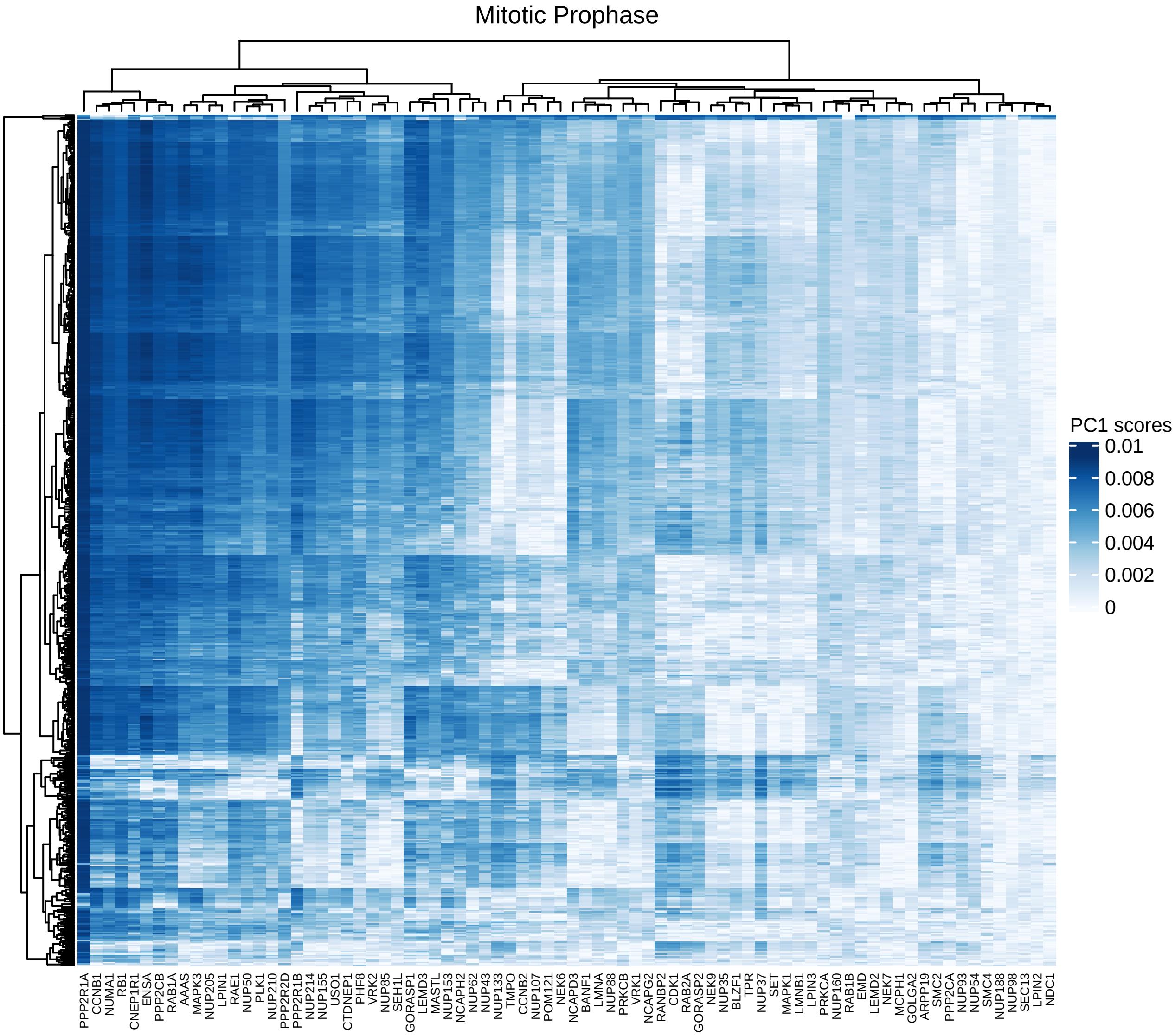

Mitotic Prophase
PC1 scores
0.01
0.008
0.006
0.004
0.002
0
PPP2R1A
CCNB1
NUMA1
RB1
CNEP1R1
ENSA
PPP2CB
RAB1A
AAAS
MAPK3
NUP205
LPIN1
RAE1
NUP50
PLK1
NUP210
PPP2R2D
PPP2R1B
NUP214
NUP155
USO1
CTDNEP1
PHF8
VRK2
NUP85
SEH1L
GORASP1
LEMD3
MASTL
NUP153
NCAPH2
NUP62
NUP43
NUP133
TMPO
CCNB2
NUP107
POM121
NEK6
NCAPD3
BANF1
LMNA
NUP88
PRKCB
VRK1
NCAPG2
RANBP2
CDK1
RAB2A
GORASP2
NEK9
NUP35
BLZF1
TPR
NUP37
SET
MAPK1
LMNB1
LPIN3
PRKCA
NUP160
RAB1B
EMD
LEMD2
NEK7
MCPH1
GOLGA2
ARPP19
SMC2
PPP2CA
NUP93
NUP54
SMC4
NUP188
NUP98
SEC13
LPIN2
NDC1

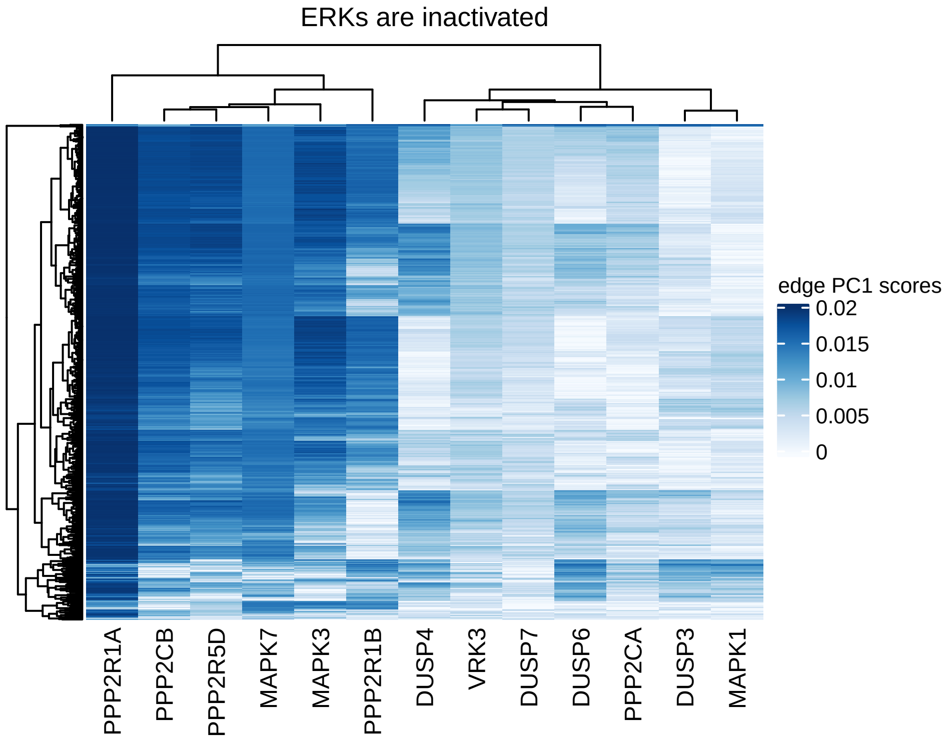

ERKs are inactivated
edge PC1 scores
0.02
0.015
0.01
0.005
0
PPP2R1A
PPP2CB
PPP2R5D
MAPK7
MAPK3
PPP2R1B
DUSP4
VRK3
DUSP7
DUSP6
PPP2CA
DUSP3
MAPK1

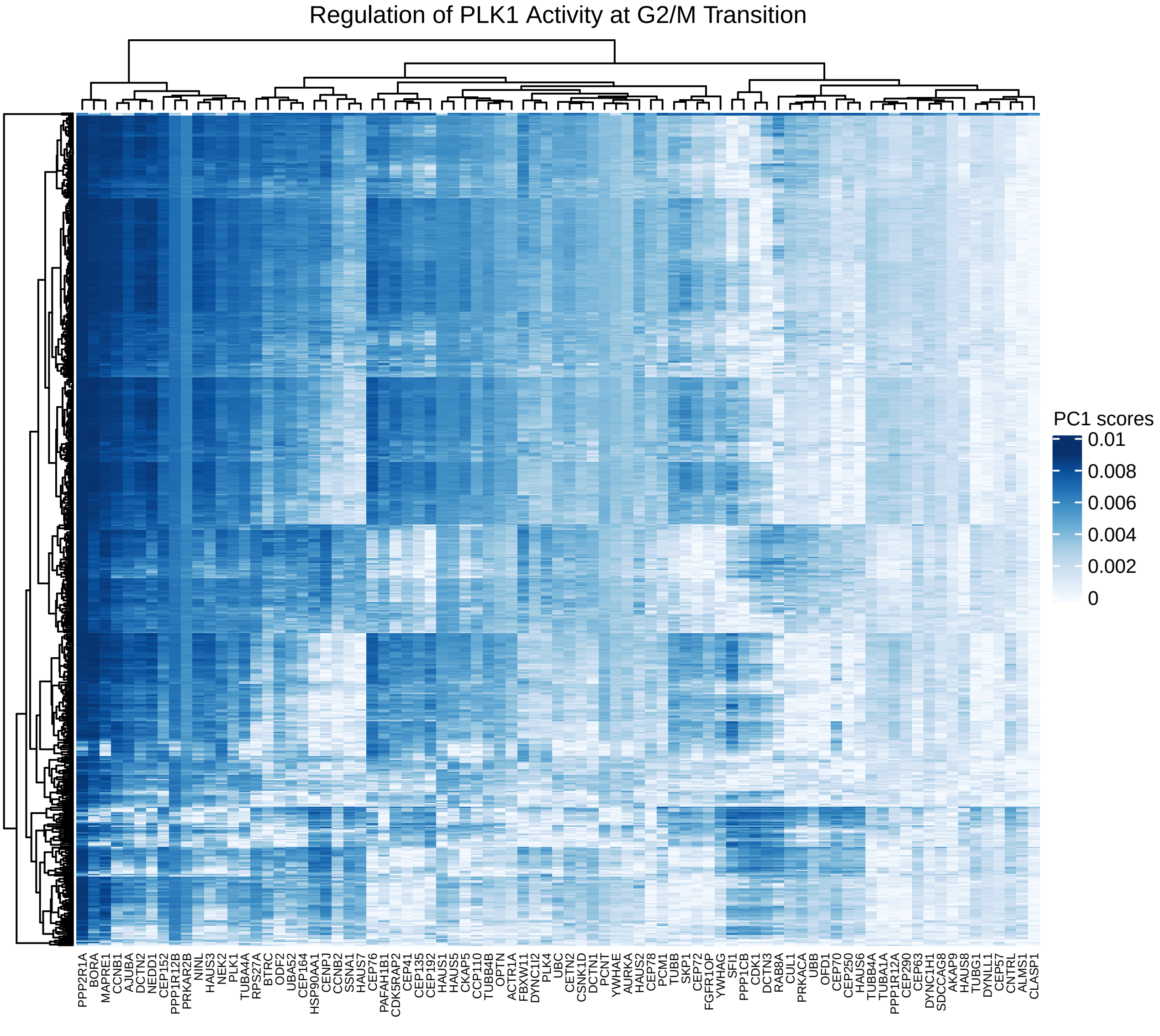
Regulation of PLK1 Activity at G2/M Transition
PC1 scores
0.01
0.008
0.006
0.004
0.002
0
PPP2R1A
BORA
MAPRE1
CCNB1
AJUBA
DCTN2
NEDD1
CEP152
PPP1R12B
PRKAR2B
NINL
HAUS3
NEK2
PLK1
TUBA4A
RPS27A
BTRC
ODF2
UBA52
CEP164
HSP90AA1
CENPJ
CCNB2
SSNA1
HAUS7
CEP76
PAFAH1B1
CDK5RAP2
CEP41
CEP135
CEP192
HAUS1
HAUS5
CKAP5
CCP110
TUBB4B
OPTN
ACTR1A
FBXW11
DYNC1I2
PLK4
UBC
CETN2
CSNK1D
DCTN1
PCNT
YWHAE
AURKA
HAUS2
CEP78
PCM1
TUBB
SKP1
CEP72
FGFR1OP
YWHAG
SFI1
PPP1CB
CDK1
DCTN3
RAB8A
CUL1
PRKACA
UBB
OFD1
CEP70
CEP250
HAUS6
TUBB4A
TUBA1A
PPP1R12A
CEP290
CEP63
DYNC1H1
SDCCAG8
AKAP9
HAUS8
TUBG1
DYNLL1
CEP57
CNTRL
ALMS1
CLASP1

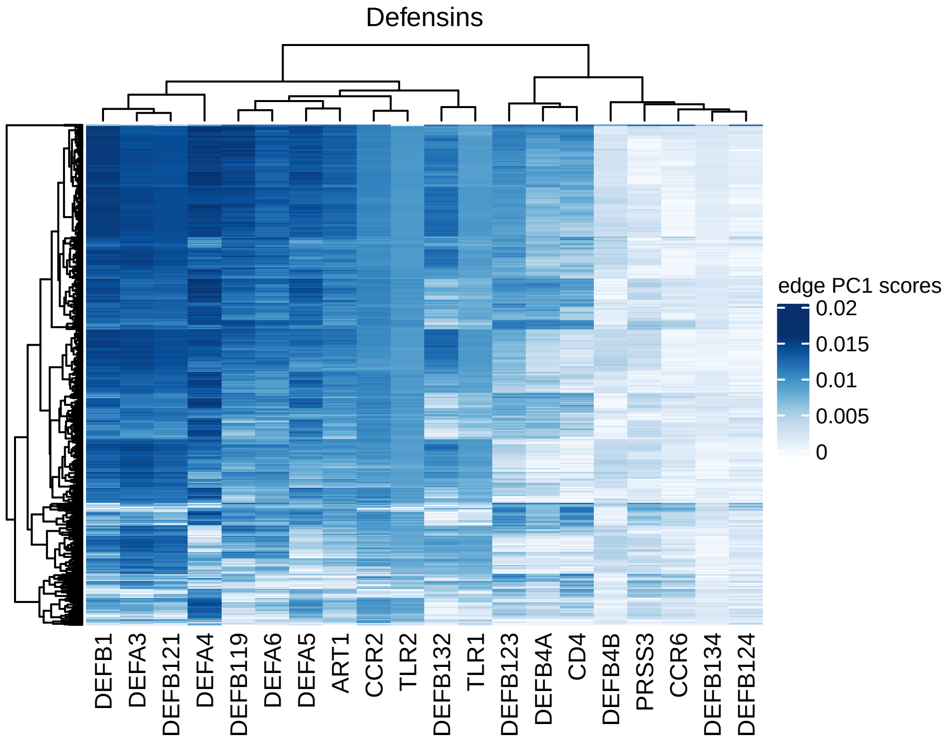
Defensins
edge PC1 scores
0.02
0.015
0.01
0.005
0
DEFB1
DEFA3
DEFB121
DEFA4
DEFB119
DEFA6
DEFA5
ART1
CCR2
TLR2
DEFB132
TLR1
DEFB123
DEFB4A
CD4
DEFB4B
PRSS3
CCR6
DEFB134
DEFB124

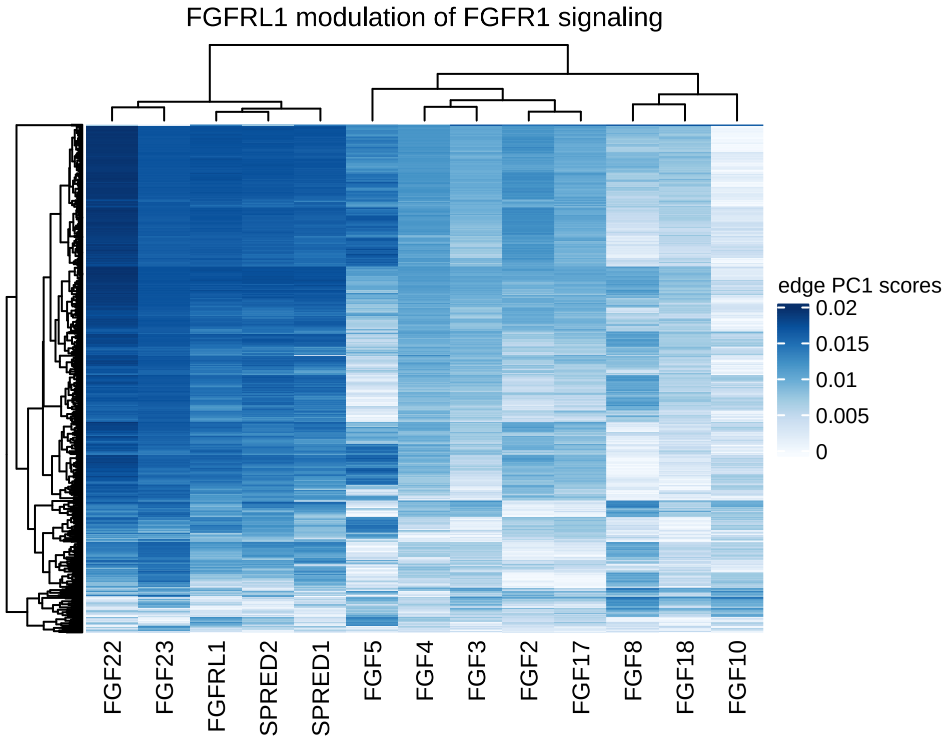

FGFRL1 modulation of FGFR1 signaling
edge PC1 scores
0.02
0.015
0.01
0.005
0
FGF22
FGF23
FGFRL1
SPRED2
SPRED1
FGF5
FGF4
FGF3
FGF2
FGF17
FGF8
FGF18
FGF10

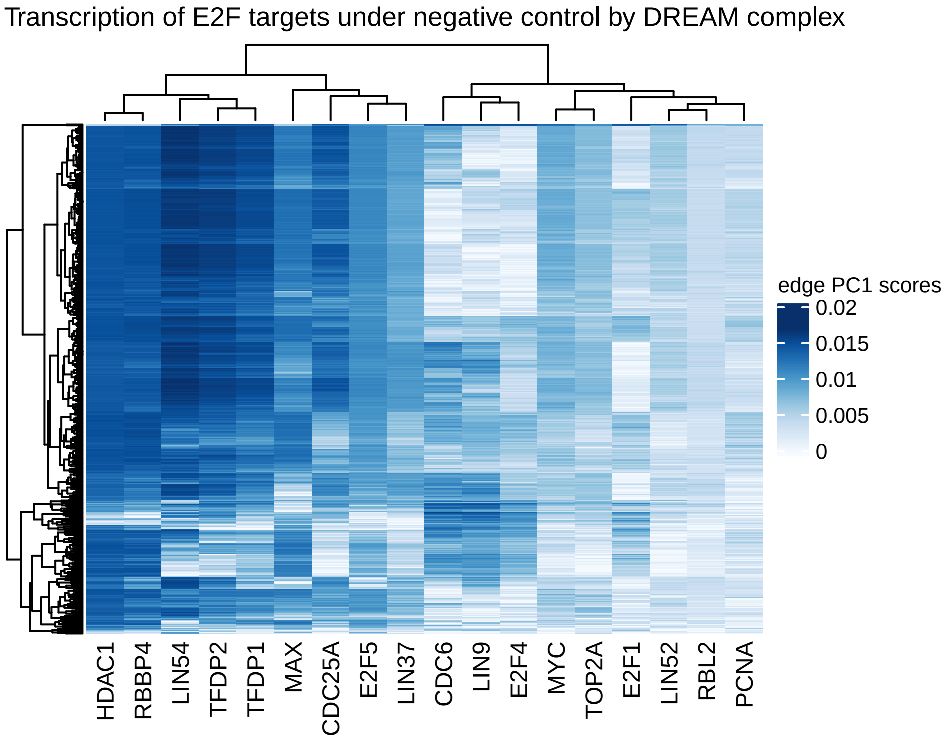
Transcription of E2F targets under negative control by DREAM complex
edge PC1 scores
0.02
0.015
0.01
0.005
0
HDAC1
RBBP4
LIN54
TFDP2
TFDP1
MAX
CDC25A
E2F5
LIN37
CDC6
LIN9
E2F4
MYC
TOP2A
E2F1
LIN52
RBL2
PCNA

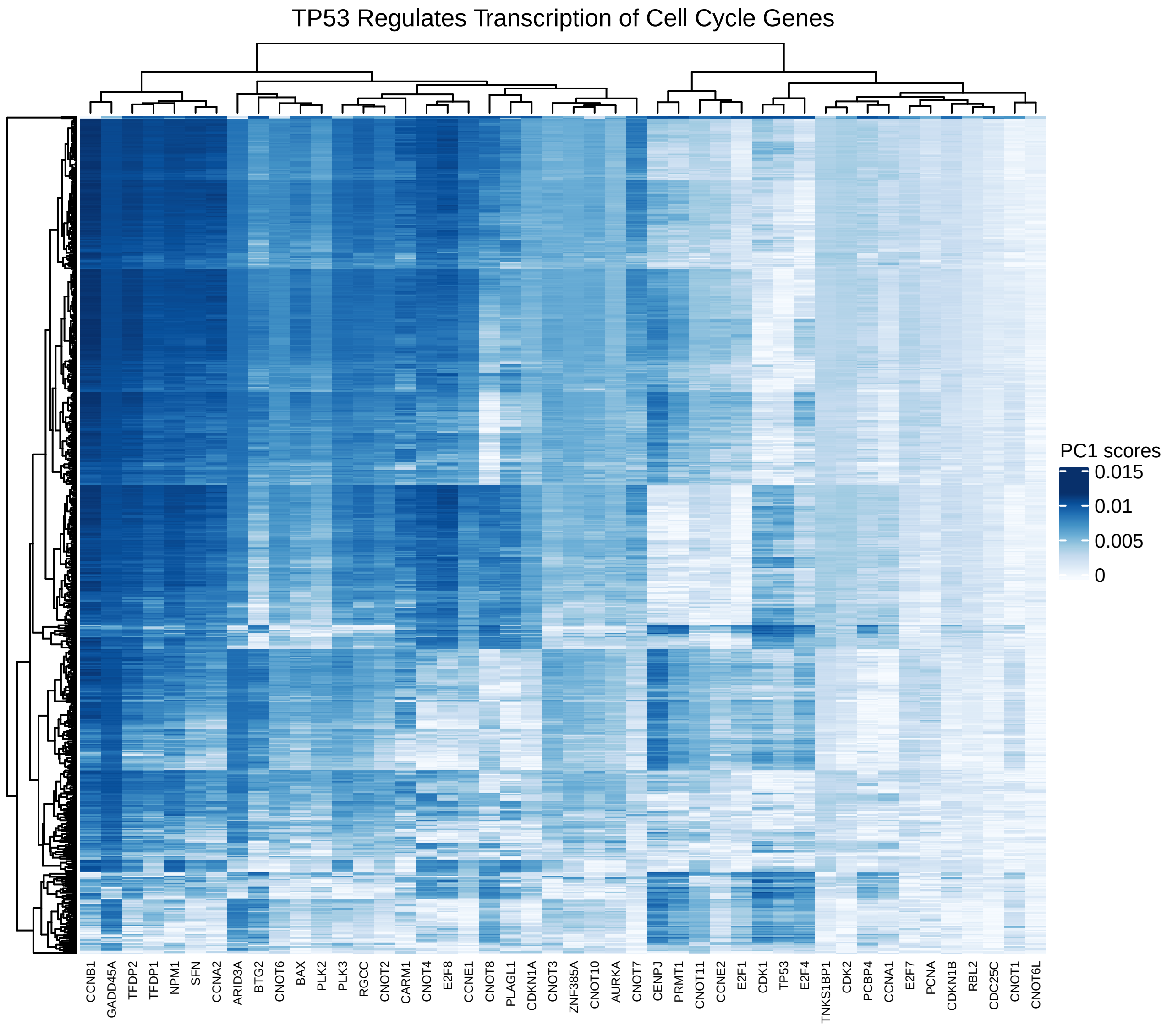
TP53 Regulates Transcription of Cell Cycle Genes
PC1 scores
0.015
0.01
0.005
0
CCNB1
GADD45A
TFDP2
TFDP1
NPM1
SFN
CCNA2
ARID3A
BTG2
CNOT6
BAX
PLK2
PLK3
RGCC
CNOT2
CARM1
CNOT4
E2F8
CCNE1
CNOT8
PLAGL1
CDKN1A
CNOT3
ZNF385A
CNOT10
AURKA
CNOT7
CENPJ
PRMT1
CNOT11
CCNE2
E2F1
CDK1
TP53
E2F4
TNKS1BP1
CDK2
PCBP4
CCNA1
E2F7
PCNA
CDKN1B
RBL2
CDC25C
CNOT1
CNOT6L

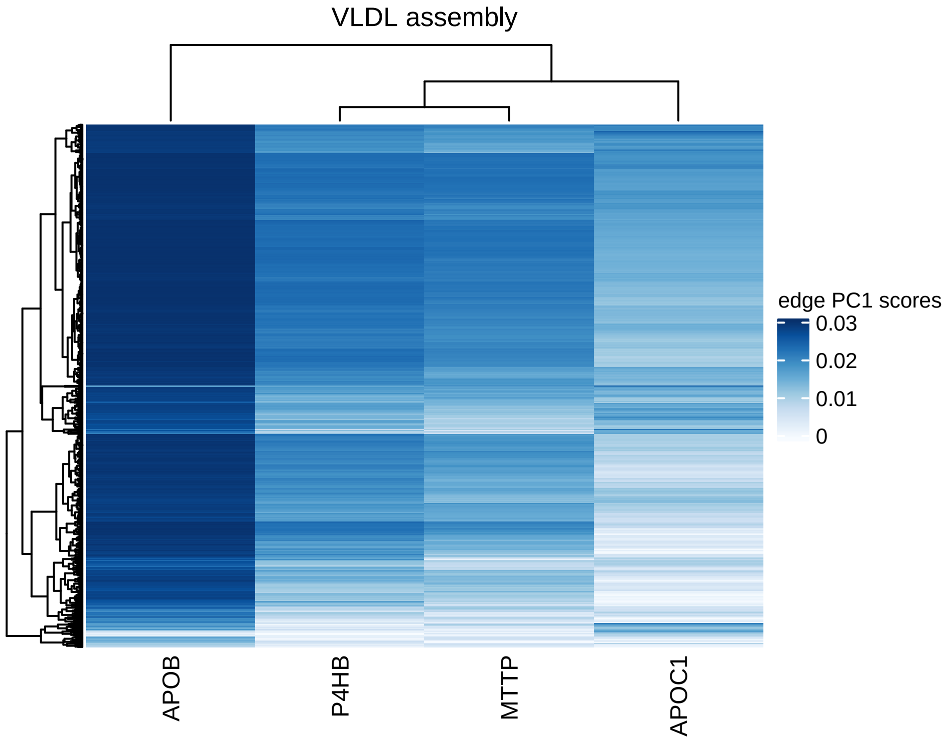
VLDL assembly
APOB
P4HB
MTTP
APOC1
edge PC1 scores
0.03
0.02
0.01
0
